# Identification of essential β-oxidation genes and corresponding metabolites for estrogen degradation by actinobacteria

**DOI:** 10.1101/2021.04.22.440886

**Authors:** Tsun-Hsien Hsiao, Tzong-Huei Lee, Meng-Rong Chuang, Po-Hsiang Wang, Menghsiao Meng, Masae Horinouchi, Toshiaki Hayashi, Yi-Lung Chen, Yin-Ru Chiang

## Abstract

Steroidal estrogens (C_18_) are contaminants receiving increasing attention due to their endocrine-disrupting activities at sub-nanomolar concentrations. Although estrogens can be eliminated through photodegradation, microbial function is critical for removing estrogens from ecosystems devoid of sunlight exposure including activated sludge, soils, and aquatic sediments. Actinobacteria were found to be key estrogen degraders in manure-contaminated soils and estuarine sediments. Previously, we used the actinobacterium *Rhodococcus* sp. strain B50 as a model microorganism to identify two oxygenase genes, *aedA* and *aedB*, involved in the activation and subsequent cleavage of the estrogenic A-ring, respectively. However, genes responsible for the downstream degradation of estrogen A/B-rings remained completely unknown. In this study, we employed tiered comparative transcriptomics, gene disruption experiments, and mass spectrometry–based metabolite profile analysis to identify estrogen catabolic genes. We observed the up-regulation of thiolase-encoding *aedF* and *aedK* in the transcriptome of strain B50 grown with estrone. Consistently, two downstream estrogenic metabolites, 5-oxo-4-norestrogenic acid (C_17_) and 2,3,4-trinorestrogenic acid (C_15_), were accumulated in *aedF-* and *aedK*-disrupted strain B50 cultures. Disruption of *fadD3* [3aα-H-4α(3’-propanoate)-7aβ-methylhexahydro-1,5-indanedione (HIP)-coenzyme A ligase gene] in strain B50 resulted in apparent HIP accumulation in estrone-fed cultures, indicating the essential role of *fadD3* in actinobacterial estrogen degradation. In addition, we detected a unique *meta*-cleavage product, 4,5-*seco*-estrogenic acid (C_18_), during actinobacterial estrogen degradation. Differentiating the estrogenic metabolite profile and degradation genes of actinobacteria and proteobacteria enables the cost-effective and time-saving identification of potential estrogen degraders in various ecosystems through liquid chromatography–mass spectrometry analysis and polymerase chain reaction–based functional assays.

## Introduction

Steroidal estrogens regulate the development of the reproductive system and secondary sex characteristics of vertebrates. The synthesis and secretion of estrogens exclusively occur in animals, particularly in vertebrates such as humans and livestock (Matsumoto et al., 1997; Tarrant et al., 2003). In the liver, estrogens undergo structural modifications (conjugation with glucuronate or sulfate) and are converted into more soluble products for subsequent excretion (Harvey and Farrier, 2011). In the environment, these conjugated estrogens are often hydrolyzed by microorganisms to form free estrogens (Koh et al., 2008). Chronic exposure to trace estrogens at subnanomolar levels can disrupt the endocrine system and sexual development in higher animals, particularly aquatic species (Belfroid et al., 1999; Baronti et al., 2000; Huang and Sedlak, 2001; Kolodziej et al., 2003; Lee et al., 2006). For example, the EC50 of 17β-estradiol (E2) that causes infertility in fathead minnows (*Pimephales promelas*) is 120 ng/L (Kramer et al., 1998). In addition, estrogens have been classified as Group 1 carcinogens by the World Health Organization (Agents classified by the *IARC Monographs*, *Volumes 1–129*). However, a recent study argued that invertebrates incapable of synthesizing estrogens are not affected by environmental estrogens (Balbi et al., 2019)

Concern is increasing about the role of estrogens as a contaminant that poses a public health challenge to municipalities due to the increasing human population and mounting demand for livestock products. Livestock manure (Hanselman et al., 2003) and municipal sewage–derived fertilizers (Lorenzen et al., 2004; Hamid and Eskicioglu, 2012) are major sources of environmental estrogens, whereas anaerobic digestion processes do not appear to alter the total estrogen concentration (<10%) in livestock manure (Noguera-Oviedo and Aga, 2016). Estrogens in sludges may be released to aquatic ecosystems through rainfall and leaching (Hanselman et al., 2003; Kolodziej et al., 2004).

Both natural estrogens [e.g., estrone (E1) and E2] and synthetic estrogens (e.g., 17α-ethynylestradiol) can be photodegraded in surface water ecosystems with a degradation half-live ranging from days to weeks (Jurgens et al., 2002; Lin and Reinhard, 2005). However, photodegradation rarely occurs in environments such as sludge, soils, and aquatic sediments that do not receive sunlight exposure. Alternatively, microbial degradation is a major mechanism for removing estrogens from these environment (Thayanukul et al., 2010; Chen et al., 2017 and 2018; Chiang et al., 2020; Wang et al., 2020). Common bacterial taxa capable of complete estrogen degradation include actinobacteria, such as *Nocardia* sp. strain E110 (*Coombe *et al*.,* 1966) and *Rhodococcus* spp. (Yoshimoto et al., 2004; Kurisu et al., 2010; Hsiao et al., 2021), and proteobacteria, such as *Novosphingobium tardaugens* (Fujii et al., 2002), *Novosphingobium* spp. (Chen et al., 2018; Wu et al., 2019; Li et al., 2021), *Sphingobium estronivorans* (Qin et al., 2020), and *Sphingomonas* spp. (Ke et al., 2007; Yu et al., 2007; Chen et al., 2017). Several possible estrogen biodegradation pathways have been proposed (Yu et al., 2013; Chiang et al., 2020; Li et al., 2021), suggesting that bacteria in different taxa likely adopt multiple strategies to degrade estrogens. The results of gene disruption experiments and enzyme characterization have indicated that proteobacteria adopt the cytochrome P450-type monooxygenase EdcB (estrone 4-hydroxylase; Ibero et al., 2020) and the type I extradiol dioxygenase OecC (4-hydroxyestrone 4,5-dioxygenase; Chen et al., 2017) to activate and cleave the estrogenic A-ring, respectively. Recently, we demonstrated that actinobacteria use a similar strategy to cleave the estrogenic A-ring with functionally homologous enzymes exhibiting a sequence identity of <40% for proteobacterial enzymes. However, essential genes involved in the downstream steps of both actinobacterial and proteobacterial degradation pathways remain unknown.

In this study, we used the actinobacterium *Rhodococcus* sp. strain B50, a soil isolate, as the model microorganism to identify downstream metabolites and catabolic genes because of its excellent efficiency in estrogen degradation and its compatibility with commercial genetic manipulation techniques. On the basis of the annotated strain B50 genome (Hsiao et al., 2021), we performed tiered comparative transcriptomics, gene disruption experiments, and metabolite profile analysis to elucidate the actinobacterial estrogen degradation pathway.

## Results

### Identification of strain B50 genes involved in estrogen degradation through comparative transcriptomic analysis

We first employed comparative transcriptomics to probe genes differentially expressed under E1-fed conditions. In addition to the degradation of estrogens such as E1 and E2, strain B50 can degrade other steroids such as testosterone and cholesterol. Therefore, we used the transcriptomes of strain B50 grown on cholesterol and testosterone as controls for the comparative transcriptomics analysis. Consistent with the observed phenotype, the strain B50 linear chromosome (GMFMDNLD 2) contains a complete set of cholesterol/androgen degradation genes in the established 9,10-*seco* pathway (Holert et al., 2016; Crowe et al., 2018) including genes involved in cholesterol uptake (*mce4* genes; GMFMDNLD_02935 to 02949), steroidal side-chain degradation (GMFMDNLD_02968 to 02992 and GMFMDNLD_03076 to 03082), androgenic A/B-rings degradation (GMFMDNLD_03002 to 03014 and GMFMDNLD _03061 to 03069), and steroidal C/D-rings degradation (GMFMDNLD _03017 to 03024 and GMFMDNLD _03033 to 03050; **Dataset S1**). Among them, we identified *fadD3* (HIP-CoA ligase gene; GMFMDNLD_03043) as being responsible for steroidal C-rings degradation.

In our previous study (Hsiao et al., 2021), we identified the *aed* gene cluster containing two oxygenase genes, *aedA* and *aedB*, that is located in the megaplasmid of strain B50. Here, we investigated whether *aed* genes are induced by estrogens. We grew strain B50 cells by using three steroid substrates, namely E1, testosterone, or cholesterol. Subsequently, we performed a comparative transcriptomic analysis to detect genes specifically up-regulated in the E1-fed culture (**Fig. 1**). As expected, *mce4* genes (GMFMDNLD_02935 to 02949) were apparently up-regulated only under cholesterol-fed conditions (**Fig. 1A**). Our data indicated that genes involved in steroidal C- and D-rings are expressed at similar levels (<4-fold difference) in all three treatments (**Fig. 1**). By contrast, the *aed* gene cluster (GMFMDNLD _05332 to 05349) was differentially up-regulated (>5-fold difference) in the E1-fed culture (**Fig. 1; Dataset S1**). In addition to GMFMDNLD _05336 and GMFMDNLD _05338 encoding AedA and AedB, respectively (Hsiao et al., 2021), in the *aed* gene cluster, we identified a putative medium-chain fatty acid:CoA ligase gene [GMFMDNLD _05341 (*aedJ*)]. Moreover, genes encoding two sets of β-oxidation enzymes, including acyl-CoA dehydrogenase [GMFMDNLD _05345 (*aedN*) and 05347 (*aedP*)], enoyl-CoA hydratase [GMFMDNLD _05333 (*aedD*), 05344 (*aedM*), and 05346 (*aedO*)], 3-hydroxyacyl-CoA dehydrogenase [GMFMDNLD _05334 (*aedE*) and 05337 (*aedG*)], and thiolase [GMFMDNLD _05335 (*aedF*) and 05342 (*aedK*)], are present in this gene cluster (**Fig. 2**). Highly similar β-oxidation genes (with a deduced amino acid sequence identity >70%) were also present in the genome of the estrogen-degrading *Rhodococcus* sp. strain DSSKP-R-001.

**FIG 1.**
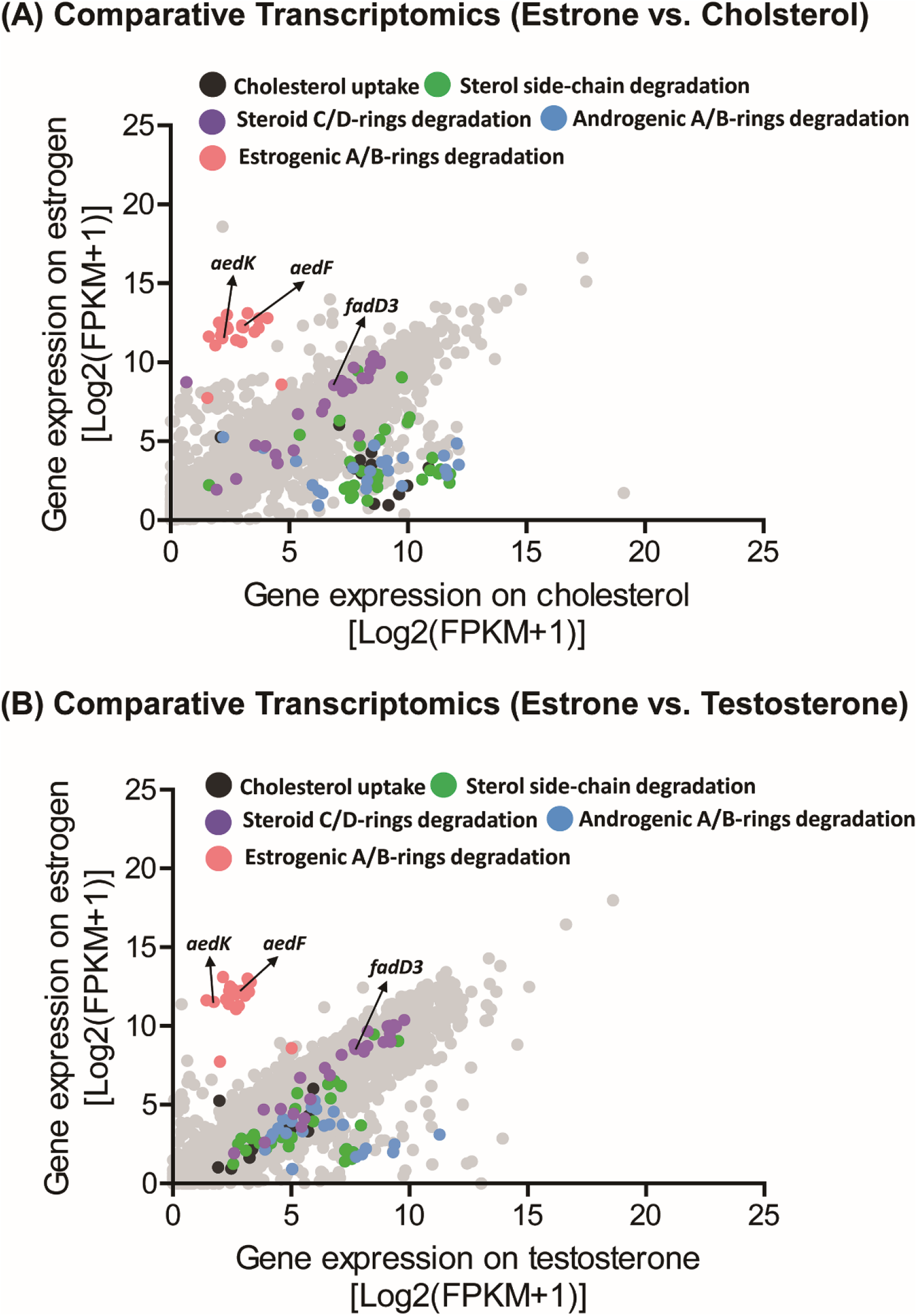
Comparative transcriptomic analyses of *Rhodococcus* sp. strain B50. (A) Global gene expression profiles (RNA-Seq) of strain B50 grown on estrone (E1) or cholesterol. (B) Global gene expression profiles of strain B50 grown on E1 or testosterone. Each spot represents a gene.

**FIG 2.**
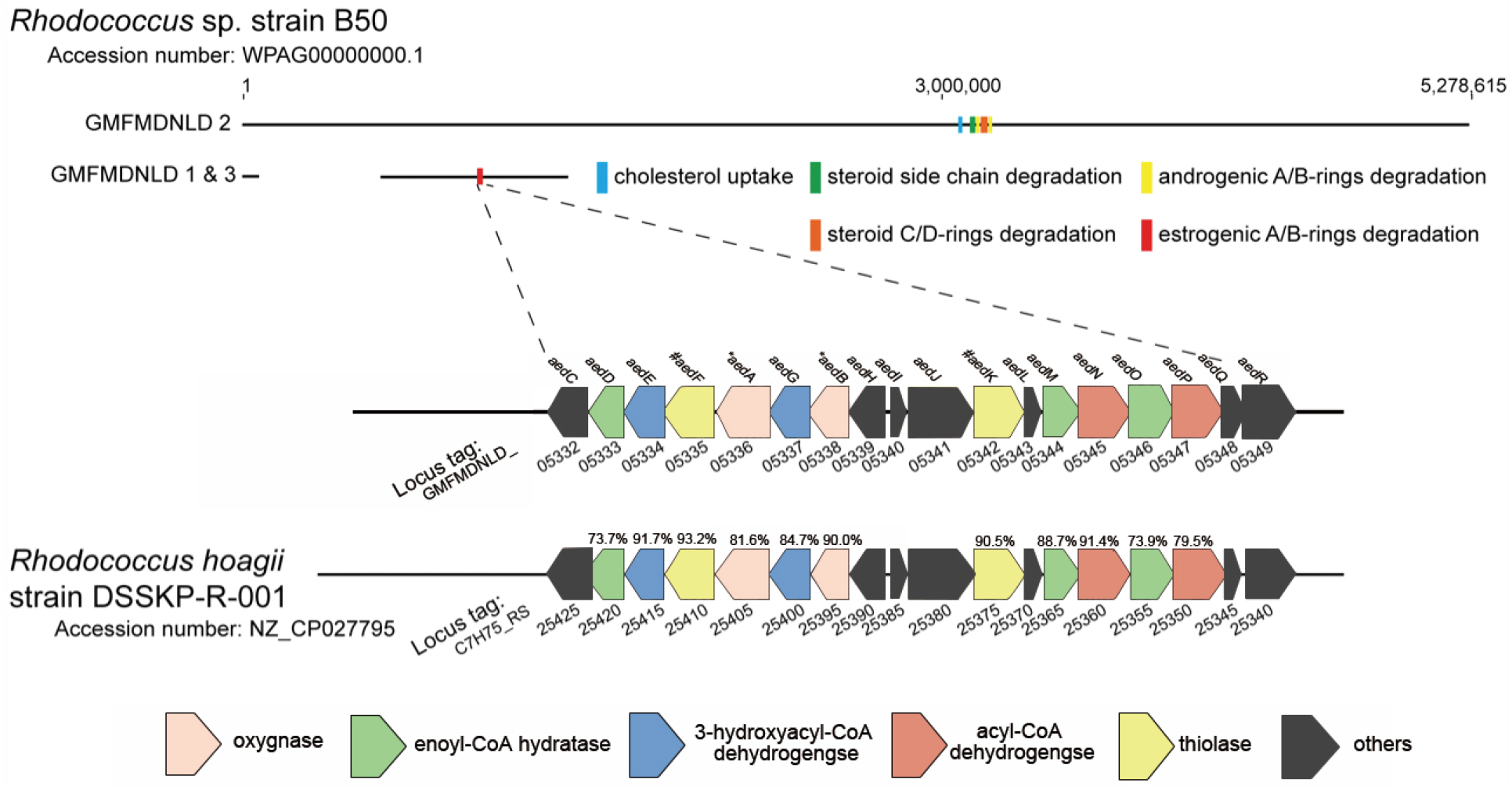
The *aed* gene cluster specific for the actinobacterial degradation of estrogenic A- and B-rings. Percentage (%) indicates the shared identity of the deduced amino acid sequences of *Rhodococcus* sp. strain DSSKP-R-001. *, the oxygenase genes *aedA* and *aed* B were characterized in our previous study (Hsiao et al., 2021).

### Functional validation of β-oxidation genes involved in estrogen biodegradation

Next, we elucidated the function of putative β-oxidation genes involved in actinobacterial estrogen degradation. Thus, we disrupted two putative thiolase genes [GMFMDNLD_05335 (*aedF*) and GMFMDNLD_05342 (*aedK*)] of strain B50 through site-directed mutagenesis [insertion of a chloramphenicol-resistance gene (Cm^R^) and *pheS*\*\* cassette] (Fig. 3A). The plasmid was transferred from *Escherichia coli* (nalidixic acid-sensitive) to strain B50 (nalidixic acid-resistant) through conjugation. Subsequently, the gene-disrupted strain B50 mutants were selected and maintained on lysogeny broth (LB) agar containing two antibiotics: chloramphenicol (25 μg/mL) and nalidixic acid (12.5 μg/mL). Polymerase chain reaction (PCR) with primers flanking *aedF* and Cm^R^ genes confirmed the successful insertion of the chloramphenicol-resistance cassette into the *aedF* gene in the mutated strain (Fig. 3B). The results of metabolite profile analysis indicated the accumulation of a major C_17_ product (**Metabolite 3**) in the E1-fed bacterial culture of an *aedF*-disrupted mutant but not in that of the wild-type strain B50 (**Fig.4**). Applying the same gene-disruption approach (insertion of the Cm^R^ and *pheS*\*\* cassette), we obtained an *aedK*-disrupted strain B50 mutant (**Fig. 3B**). We observed the accumulation of a C_15_ metabolite (**Metabolite 7**) in *aedK*-disrupted strain B50 mutant cultures incubated with E1 (**Fig. 4**).

**FIG 3.**
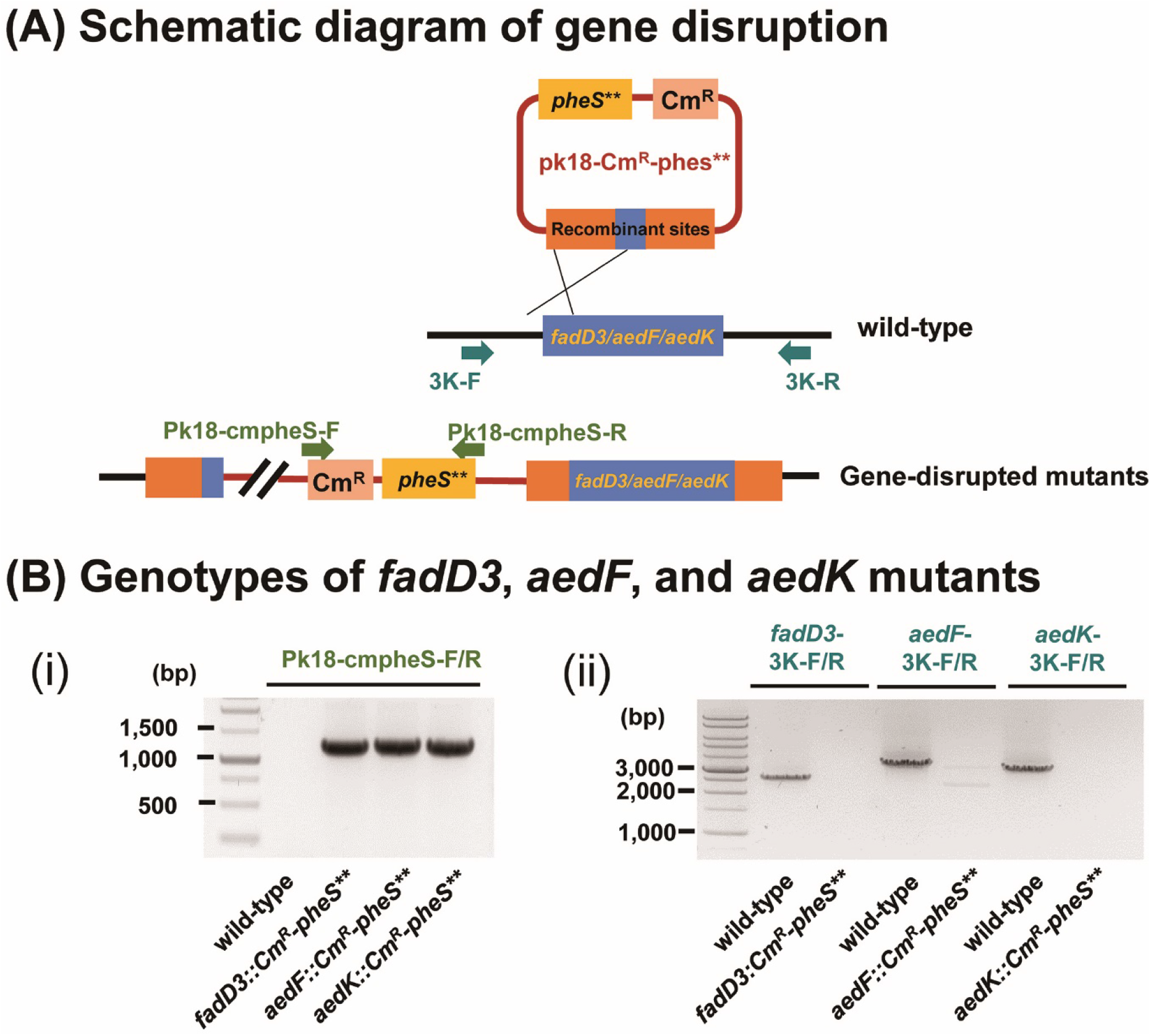
Genotype examinations of strain B50 mutants. (**A**) Schematic of homologous recombination-mediated gene disruption. (**B**) Genotype examinations of gene-disrupted mutants. Agarose gel electrophoresis indicated (**Bi**) the insertion of a chloramphenicol-resistant gene (Cm^R^) and *pheS*\*\* cassette into target genes and (**Bii**) the gene disruption of strain B50 mutants. The wild-type strain B50 was also tested for comparison.

**FIG 4.**
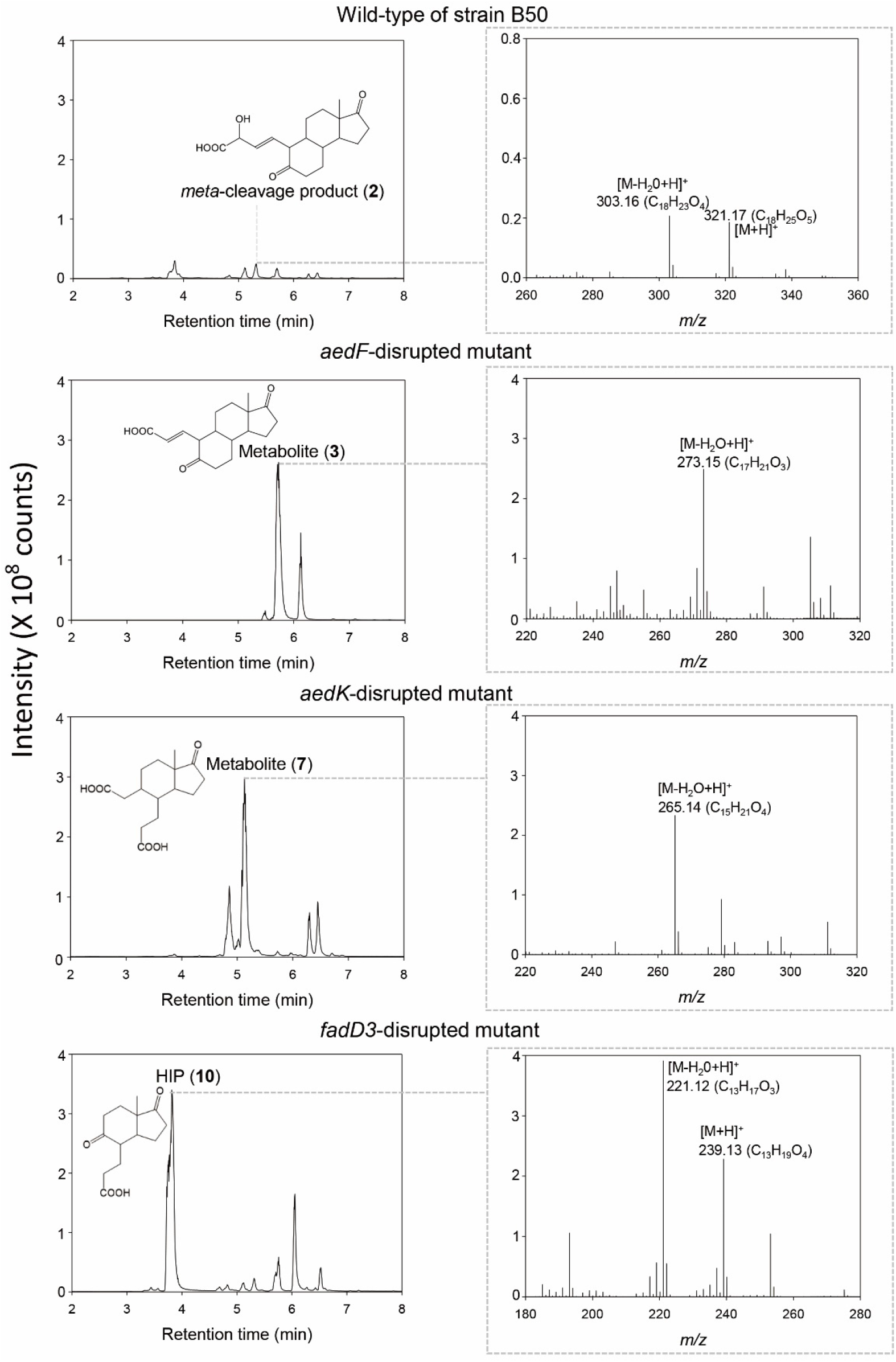
Validation of the phenotypes of strain B50 mutants. UPLC−APCI−HRMS analysis indicated the apparent accumulation of C_17_ (Metabolite **3**), C_15_ (Metabolite **7**), and C13 (HIP) metabolites in the E1-fed cultures of *aedF*-, *aedK*-, and *fadD3*-disrupted mutants, respectively. The major estrogenic metabolites observed in the wild-type strain B50 culture include the C_18_ *meta*-cleavage product (Metabolite **2**).

Studies have identified 3aα-H-4α(3’-propanoate)-7aβ-methylhexahydro-1,5-indanedione (HIP) as a possible estrogenic metabolite for proteobacteria (Wu et al., 2017) and actinobacteria (Hsiao et al., 2021). However, whether *fadD3* (HIP-CoA ligase gene; GMFMDNLD_03043) is essential for estrogen degradation remain unclear. Therefore, we disrupted *fadD3* with the Cm^R^ and *pheS*\*\* cassette (**Fig. 3B**) and compare the E1 degradation of the wild type and *fadD3*-disrupted mutant of strain B50. The *fadD3*-disrupted mutant transformed E1 into HIP and apparently accumulated HIP in bacterial cultures (**Fig. 4**). By contrast, the wild-type strain B50 completely degraded E1 (1 mM) within 8 h and did not apparently accumulated HIP (<0.1 mM) as an end product in bacterial cultures.

### Structural elucidation of novel estrogen metabolites through nuclear magnetic resonance spectroscopy

We elucidated the structures of estrogenic metabolites accumulated in the wild-type strain B50 cultures [e.g., 4-hydroxyestrone, pyridinestrone acid (PEA), Metabolite 2, and HIP] and those accumulated in the bacterial cultures of strain B50 mutants including Metabolites 3 and 7(Table 1). Subsequently, we identified the structures of three novel estrogen-derived metabolites (Fig. 5). Among them, a high-performance liquid chromatography (HPLC)-purified compound (Metabolite 3), obtained as a colorless oil, was assigned a molecular formula of C_17_H_22_O_4_ by a quasi-molecular adduct [M + H]^+^ at *m/z* 291.16 (calculated as 291.1596 for C_17_H_23_O_4_) in the positive ion mode of ultraperformance liquid chromatography–electrospray ionization–high-resolution mass spectrometry (UPLC–ESI–HRMS; Fig. S1). When we compared the ^1^H- and ^13^C-NMR data of Metabolite 3 with those of the previously reported 4-norestrogenic acid (Wu et al., 2019), the NMR data of Metabolite 3 were almost identical with those of 4-norestrogenic acid except that a carbonyl signal at δ_H_ 3.35 (H-5)/δ_C_ 73.8 in 4-norestrogenic acid disappeared and an additional ketone signal at δ_C_ 211.7 (C-5) was observed in Metabolite 3(Table 2; see Fig. S3 for original NMR spectra), indicating that the hydroxy group at C-5 of 4-norestrogenic acid was substituted by a carbonyl in Metabolite 3. In addition, this change was evidenced by the distinctive downfield shifts of the ^13^C-NMR data of C-1, C-6, C-7, C-9, and C-10 of Metabolite 3, upfield shifts of the ^13^C-NMR data of its C-2 and C-3, and key HMBC cross-peaks of δ_H_ 2.55 and 2.42 (H_2_-6)/δ_C_ 211.7 (C-5), δ_H_ 3.00 (H-10)/δ_C_ 211.7 (C-5), and δ_H_ 6.78 (H-1)/δ_C_ 211.7 (C-5) (**Figs. 5** and **S6**). The configuration of the double bond at C-1 and C-2 of Metabolite **3** was established to be an *E* form based on the larger coupling constant (*J*_H-1/H-2_ = 15.6 Hz). Thus, the structure of Metabolite **3** was deduced and is shown in **Fig. 5** —it it was named 5-oxo-4-norestrogenic acid.

**FIG 5.**
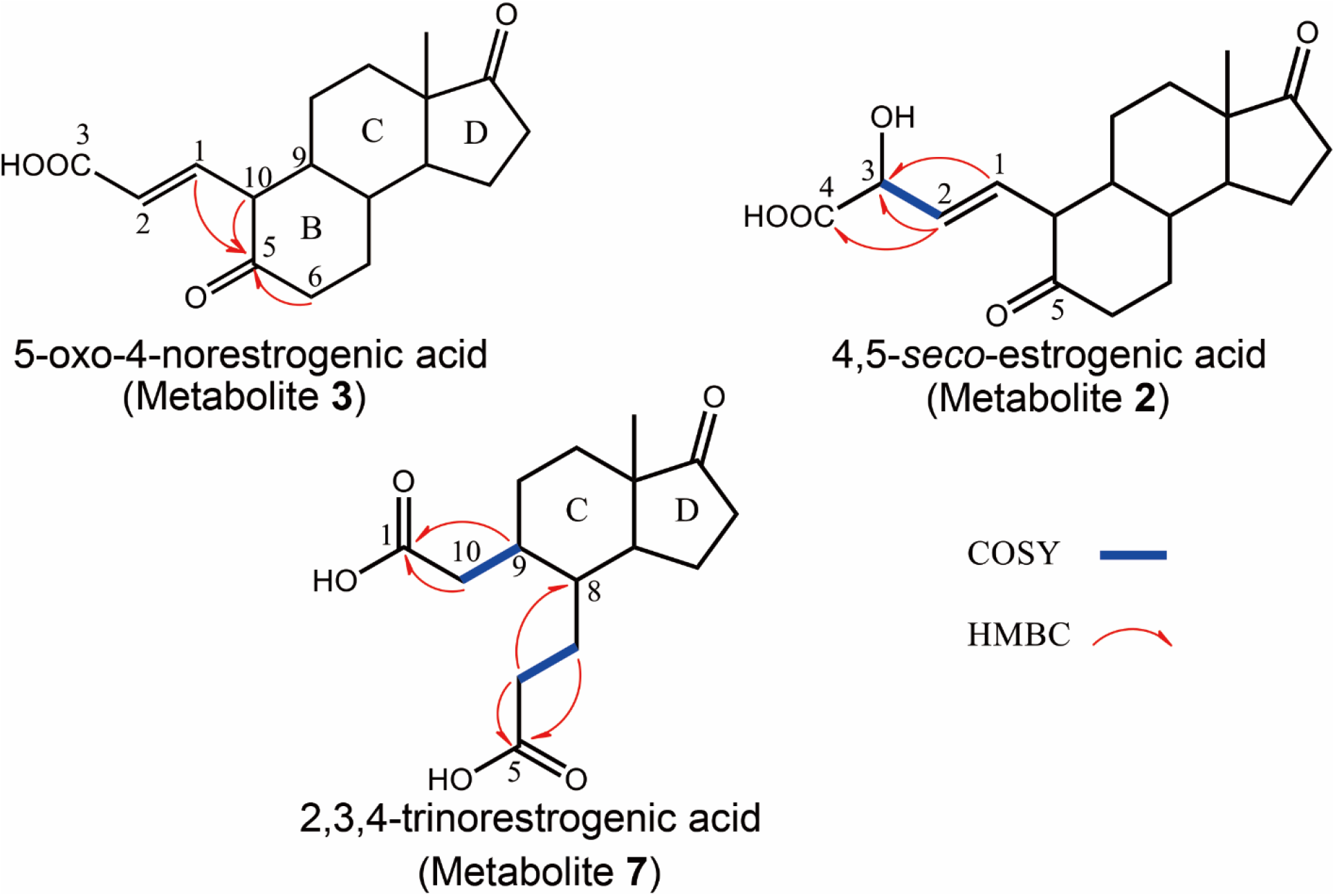
Key COSY and HMBC correlations in the two-dimensional NMR data of three HPLC-purified metabolites produced by strain B50. Original COSY and HMBC spectra of Metabolites **2**, **3**, and **7** are provided in **Figs. S5**, **S6**, and **S7**, respectively.

**Table 1.** UPLC-HRMS analysis of metabolites involved in estrogen degradation by strain B50. Estrogen metabolites newly identified in this study are boldfaced.

| Compound ID | UPLC behavior (RT<sup>a</sup>, min) | Molecular formula/ (predicted molecular mass)<sup>b</sup> | Dominant ion peaks | Identification of product ions | Mode observed |
| --- | --- | --- | --- | --- | --- |
| E1 | 8.13 | C<sub>18</sub>H<sub>22</sub>O<sub>2</sub><br>270.16 | 253.16<br>271.17 | [M-H<sub>2</sub>O+H]<sup>+</sup><br>[M+H]<sup>+</sup> | ESI and APCI<br>ESI and APCI |
| 4-hydroxyestrone | 7.40 | C<sub>18</sub>H<sub>22</sub>O<sub>3</sub><br>286.16 | 269.16<br>287.15<br>309.15 | [M-H<sub>2</sub>O+H]<sup>+</sup><br>[M+H]<sup>+</sup><br>[M+Na]<sup>+</sup> | APCI<br>ESI and APCI<br>ESI |
| PEA* | 4.02 | C<sub>18</sub>H<sub>21</sub>O<sub>3</sub>N<br>299.15 | 282.17<br>300.16<br>322.15 | [M-H<sub>2</sub>O+H]<sup>+</sup><br>[M+H]<sup>+</sup><br>[M+Na]<sup>+</sup> | ESI<br>ESI and APCI<br>ESI |
| **4,5-*seco*-estrogenic acid**<br>(Metabolite 2) | 5.24 | C<sub>18</sub>H<sub>24</sub>O<sub>5</sub><br>320.17 | 303.16<br>321.17<br>343.15 | [M-H<sub>2</sub>O+H]<sup>+</sup><br>[M+H]<sup>+</sup><br>[M+Na]<sup>+</sup> | ESI and APCI<br>ESI and APCI<br>ESI |
| **5-oxo-4-norestrogonic acid**<br>(Metabolite 3) | 5.82 | C<sub>17</sub>H<sub>22</sub>O<sub>4</sub><br>290.16 | 273.15<br>291.16<br>313.14 | [M-H<sub>2</sub>O+H]<sup>+</sup><br>[M+H]<sup>+</sup><br>[M+Na]<sup>+</sup> | ESI and APCI<br>ESI and APCI<br>ESI |
| **2,3,4-trinorestrogonic acid**<br>(Metabolite 7) | 5.08 | C<sub>15</sub>H<sub>22</sub>O<sub>5</sub><br>282.15 | 247.13<br>265.14<br>283.15 | [M-2H<sub>2</sub>O+H]<sup>+</sup><br>[M-H<sub>2</sub>O+H]<sup>+</sup><br>[M+H]<sup>+</sup> | ESI and APCI<br>ESI and APCI<br>ESI and APCI |
| HIP | 3.81 | C<sub>13</sub>H<sub>18</sub>O<sub>4</sub><br>238.12 | 221.12<br>239.13<br>261.11 | [M-H<sub>2</sub>O+H]<sup>+</sup><br>[M+H]<sup>+</sup><br>[M+Na]<sup>+</sup> | ESI and APCI<br>ESI and APCI<br>ESI |

<sup>a</sup>RT, retention time. <sup>b</sup>The predicated molecular mass was calculated using the atom masses of <sup>12</sup>C (12.00), <sup>16</sup>O (15.99), and <sup>1</sup>H (1.01). \*PEA is a dead-end product.

**Table 2.** ^1^H-(600 MHz) and ^13^C-NMR (150 MHz) spectral data o f 5-oxo-4-norestrogenic acid (Metabolite **3**), 4,5-*seco*-estrogenic acid (Metabolite **2**), and 2,3,4-trinorestrogenic acid (Metabolite **7**).

| Positions | Metabolite **3** |  | Metabolite **2** |  | Metabolite **7** |  |
| --- | --- | --- | --- | --- | --- | --- |
| | $^1\text{H}^{a, b}$ | $^{13}\text{C}^a$ | $^1\text{H}^{a, b}$ | $^{13}\text{C}^a$ | $^1\text{H}^{a, b}$ | $^{13}\text{C}^a$ |
| 1 | 6.78 dd (9.6, 15.6) | 147.1 | 5.73 dd (10.2, 16.2) | 131.5 |  | 177.1 |
| 2 | 5.79 d (15.6) | 126.2 | 5.56 dd (6.3, 16.2) | 132.5 |  |  |
| 3 |  | 169.4 | 4.63 d (6.3) | 72.8 |  |  |
| 4 |  |  |  | 176.1 |  |  |
| 5 |  | 211.7 |  | 213.3 |  | 177.8 |
| 6 | 2.55 m | 42.2 | 2.54 m | 42.3 | 2.33 m | 30.5 |
|  | 2.42 m |  | 2.41 m |  |  |  |
| 7 | 2.20 m | 31.5 | 2.18 m | 31.7 | 1.82 m | 25.3 |
|  | 1.44 m |  | 1.43 m |  |  |  |
| 8 | 1.90 m | 40.4 | 1.83 m | 40.6 | 1.60<sup>d</sup> | 39.9 |
| 9 | 1.41 m | 49.9 | 1.28 m | 50.5 | 1.60<sup>d</sup> | 39.2 |
| 10 | 3.18 t (10.2) | 59.9 | 3.00 t (10.2) | 60.2 | 2.60 m | 39.0 |
|  |  |  |  |  | 2.13<sup>c</sup> |  |
| 11 | 1.60 m | 28.7 | 1.73 m | 28.6 | 1.76 m | 29.0 |
|  | 1.40 m |  | 1.34 m |  | 1.48 m |  |
| 12 | 1.75 m | 32.3 | 1.72 m | 32.4 | 1.69 m | 32.6 |
|  | 1.28 m |  | 1.26 m |  | 1.28 m |  |
| 13 |  | 49.4 |  | 49.5 |  | 49.5 |
| 14 | 1.47 m | 51.2 | 1.42 m | 51.4 | 1.53 m | 50.0 |
| 15 | 2.02 m | 22.8 | 2.04 m | 22.8 | 2.02 m | 23.5 |
|  | 1.67 m |  | 1.70 m |  | 1.69 m |  |
| 16 | 2.48 dd (8.4, 19.2) | 36.7 | 2.48 dd (8.4, 19.2) | 36.8 | 2.46 dd (9.0, 19.2) | 36.7 |
|  | 2.13 dd (9.6, 19.2) |  | 2.10 dd (9.0, 19.2) |  | 2.13<sup>c</sup> |  |
| 17 |  | 223.1 |  | 223.3 |  | 223.7 |
| 18 | 0.98 s | 14.4 | 0.98 s | 14.5 | 0.90 s | 14.1 |

<sup>a</sup>Measured in methanol-*d*<sub>4</sub>; <sup>b</sup> $\delta$ in ppm, mult. (*J* in Hz). <sup>c,d</sup> Signals were overlapped and picked up from HMBC or HSQC experiments. The original NMR spectra of Metabolites **2**, **3**, and **7** are shown in **Figs. S2, S3, and S4**, respectively.

The NMR data of Metabolite **2** (**Table 2;** see **Fig. S2** for original NMR spectra) were consistent with those of Metabolite **3** except an additional carbinol signal at δ_H_ 4.63 (H-3)/δ_C_ 72.8 (C-3) was observed in Metabolite **2**. Key COSY of δ_H_ 4.63 (H-3)/δ_H_ 5.56 (H-2) accompanied with key cross-peaks of δ_H_ 5.73 (H-1)/δ_C_ 72.8 (C-3), δ_H_ 5.56 (H-2)/δ_C_ 72.8 (C-3), and δ_H_ 5.56 (H-2)/δ_C_ 176.1 (C-4) in the HMBC spectrum (**Figs. 5** and **S5**) confirmed the additional carbinol was attached at C-3. The assignments matched with the molecular formula of compound **2**, C_18_H_24_O_5_, as interpreted from a quasi-molecular adduct [M + H]^+^ at *m/z* 321.17 (calculated as 321.1702 for C_18_H_25_O_5_) in the positive ion mode of UPLC–ESI–HRMS (**Fig. S1**). The configuration of the double bond at C-1 and C-2 of Metabolite **2** was established to be an *E* form based on the larger coupling constant (*J*_H-1/H-2_ = 16.2 Hz). The structure of Metabolite **2** is shown in **Fig. 5**, and it was named 4,5-*seco*-estrogenic acid.

The NMR data of Metabolite **7** (**Table 2;** see **Fig. S4** for original NMR spectra) coincided well with those of Metabolite **3** except the notable upfield shifts of C-8 and C-9 of Metabolite **7**, indicating conspicuous changes on the B-ring of Metabolite **7**. Key COSY of δ_H_ 2.33 (H_2_-6)/δH 1.82 (H_2_-7) and δ_H_ 1.60 (H-9)/δ_H_ 2.13 and 2.60 (H_2_-10) together with key cross-peaks of δ_H_ 2.33 (H_2_-6)/δ_C_ 177.8 (C-5), δ_H_ 2.33 (H_2_-6)/δ_C_ 39.9 (C-8), δ_H_ 1.82 (H_2_-7)/δ_C_ 177.8 (C-5), δ_H_ 1.60 (H-9)/δ_C_ 177.1 (C-1), and δ_H_ 2.13 and 2.60 (H_2_-10)/δ_C_ 177.1 (C-1) in the HMBC spectrum (**Figs. 5** and **S7**) corroborated that the B-ring of Metabolite **7** was opened and with two terminal carboxylic acids at C-1 and C-5. After being further confirmed by a quasi-molecular adduct [M + H]^+^ at *m/z* 283.15 (calculated as 283.1545 for C_15_H_23_O_5_) from the UPLC–ESI–HRMS data analysis (**Fig. S1**), the structure of Metabolite **7** was elucidated, as shown in **Fig. 5**, and it was named 2,3,4-trinorestrogenic acid.

## Discussion

### Establishment of an estrogen degradation pathway in actinobacteria

In a previous study, we identified initial metabolites (namely 4-hydroxyestrone and PEA) along with two oxygenase genes (namely *aedA* and *aedB*) in strain B50 (Hsiao et al., 2021). Moreover, the findings of comparative genomic analysis indicated that this *aed* gene cluster is present in estrogen-degrading *Rhodococcus* strains. In the present study, the findings of comparative transcriptomic analysis demonstrated that *aed* genes are differentially induced by E1 but not by testosterone or cholesterol. Subsequently, we demonstrated the essential role of three genes involved in the β-oxidation of estrogenic A/B-rings including *aedF*, *aedK*, and *fadD3*. These mutants enabled the accumulation of new estrogenic metabolites in bacterial cultures. Together, an estrogen degradation pathway in actinobacteria was established (Fig. 6). The complete set of the estrogenic metabolite profile and the degradation genes of both actinobacteria and proteobacteria may allow for the facile monitoring of estrogen biodegradation in various ecosystems through liquid chromatography–mass spectrometry analysis and PCR-based functional assays.

**FIG 6.**
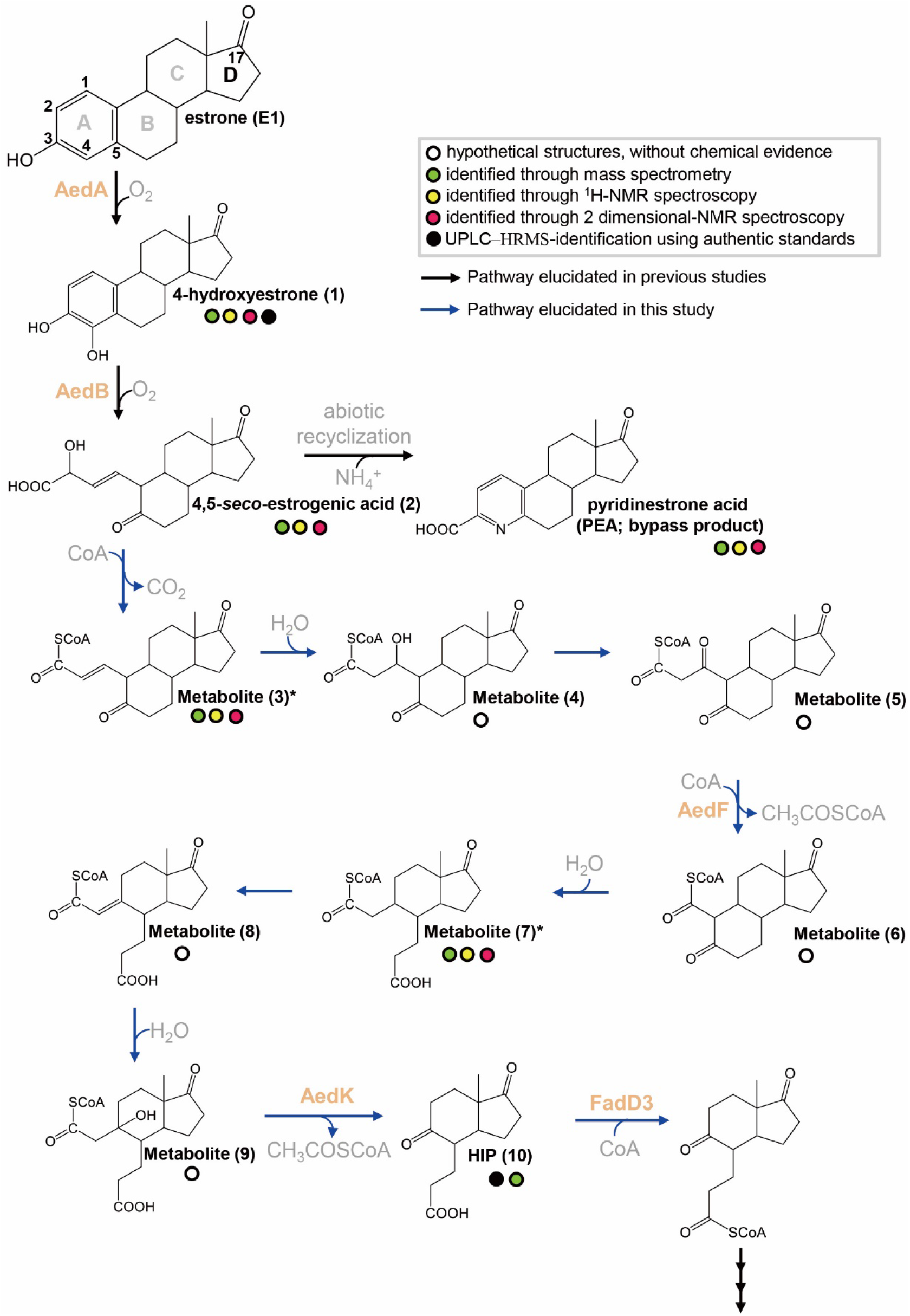
The proposed estrogen degradation pathway of actinobacteria. *, the deconjugated structures have been purified and elucidated in this study. *AedF*, *AedK*, and *fadD3* have been functionally characterized in the study.

Identification of essential β-oxidation genes and corresponding metabolites for estrogen degradation enabled the establishment of a complete degradation pathway in actinobacteria. Briefly, in the proposed pathway, the C-4 of E1 is first hydroxylated by estrone 4-hydroxylase AedA, and the resulting catecholic A-ring is cleaved through *meta*-cleavage by 4-hydroxyestrone 4,5-dioxygenase AedB. In the presence of ammonium, the *meta*-cleavage product may undergo abiotic recyclization to produce the nitrogen-containing metabolite PEA (Hsiao et al., 2021). In addition, the dead-end product PEA was detected during estrogen degradation by proteobacteria (Chen et al., 2018; Ibero et al., 2019a, 2019b, and 2020). In proteobacteria, only a minor part (approximately 2%) of the *meta*-cleavage product is abiotically transformed into PEA (Wu et al., 2019). Actinobacteria produced even less PEA (<0.5%) during E1 degradation (Hsiao et al., 2021), suggesting that the most *meta*-cleavage product was further degraded.

### Biochemical mechanisms and corresponding genes involved in estrogenic A- and B-rings degradation

Subsequent reactions may include the formation of CoA-esters through putative CoA-ligase (*aedJ*; GMFMDNLD_05341), followed the AedF-mediated removal of the C-2 and C-3 of CoA-esters through the first cycle of thiolytic β-oxidation (Fig. 6). CoA is an essential cofactor in numerous biosynthetic and energy-yielding metabolic pathways. When CoA is required in other metabolic pathways, CoA-esters in the steroid degradation pathways can be deconjugated (Takamura and Nomura, 1988; Lin et al., 2015; Wu et al., 2019). The disruption of thiolase genes thus resulted in the production and excretion of deconjugated metabolites such as Metabolites 3 and 7in E1-fed cultures.

Thus far, the mechanism operating in the actinobacterial estrogenic B-ring cleavage remains unclear. Proteobacteria degrade the estrogenic B-ring through hydrolysis (Wu et al., 2019; Ibero et al., 2020), and a similar hydrolytic ring cleavage mechanism has been demonstrated in the degradation of cyclohexanecarboxylic acid by the alphaproteobacterium *Rhodopseudomonas palustris* (Pelletier and Harwood, 1998 and 2000). Therefore, the estrogenic B-ring is likely cleaved by strain B50 through a similar hydrolytic cleavage. However, a *badI*-like gene has not been identified in the strain B50 genome, likely due to a low sequence similarity between actinobacterial and proteobacterial genes.

A highlight of the present study is the NMR identification of a C_15_ estrogenic metabolite 2,3,4-trinorestrogenic acid from *aedK*-disrupted strain B50 cultures, which validated the function of *aedK* and the proposed catabolic pathway involved in estrogenic B-ring degradation. After the hydrolytic cleavage of the estrogenic B-ring, the AedK-mediated removal of C-1 and C-10 at the CoA-ester of Metabolite **7** through the second cycle of aldolytic β-oxidation yielded HIP (**Fig. 6**). In the linear chromosome of strain B50, we identified a typical gene cluster specific for actinobacterial HIP degradation (Bergstrand et al., 2016; Crowe et al., 2018). The disruption of *fadD3* resulted in apparent HIP accumulation in E1-fed strain B50 cultures, indicating the essential role of this gene cluster in estrogenic C- and D-rings degradation.

### Activation mechanisms of the *meta*-cleavage product likely differ between actinobacteria and proteobacteria

Using strain B50 and *Sphingomonas* sp. strain KC8 as the model actinobacterium and proteobacterium, respectively, our data revealed that the actinobacterial estrogen degradation pathway is highly similar to the proteobacterial pathway (Fig. 6; Chen et al., 2017; Wu et al., 2019; Ibero et al., 2020); however, their metabolite profiles appear to be distinguishable. For example, estrogenic metabolites from actinobacteria have a typical 5-oxo group, whereas estrogenic metabolites in proteobacteria have a typical hydroxyl group at their C-5. Moreover, actinobacteria and proteobacteria seem to employ different biochemical mechanisms for estrogenic A-ring cleavage. The A-ring-cleaved metabolite in strain KC8 possesses three unsaturated double bonds at C-1, C-3, and C-5, which readily undergoes abiotic recyclization to produce PEA in the presence of ammonium, whereas strain B50 adopts an extradiol dioxygenase AedB to produce 4,5-*seco*-estrogenic acid, which has only two double bonds at C-1 and C-5 and is less likely recyclized.

Actinobacteria and proteobacteria appear to adopt different strategies to oxidize the A-ring-cleaved product (**Figure 6**). *Sphingomonas* sp. strain KC8 depends on the 2-oxoacid oxidoreductase EdcC, a member of the indolepyruvate ferredoxin oxidoreductase family, to produce 4-norestrogen-5(10)-en-3-oyl-CoA through a single oxidative decarboxylation step (Wu et al., 2019; Ibero et al., 2020). The homologous 2-oxoacid oxidoreductase gene was absent in the genome of strain B50 and any other estrogen-degrading actinobacteria. In strain B50, a putative 2-hydroxyacid dehydrogenase gene (GMFMDNLD_05337; *aedG*), a decarboxylase gene (GMFMDNLD_05339; *aedH*), and a CoA-ligase gene (GMFMDNLD_05341; *aedJ*) on the megaplasmid likely serve to add a CoA onto 4,5-*seco*-estrogenic acid (namely Metabolite **2**) through three different steps. Nevertheless, the functions of these genes remain to be characterized.

## Experimental procedures

### Chemicals

E1, E2, E3, 4-hydroxyestrone, 17α-ethynylestradiol, and HIP, also known as 3-(7a-methyl-1,5-dioxooctahydro-1H-inden-4-yl) propanoic acid, were purchased from Sigma-Aldrich (St. Louis, Missouri, USA). PEA was prepared as described by Chen et al., 2017. The [3,4C-^13^C]E1 (99%) was purchased from Cambridge Isotope Laboratories (Tewksbury, Massachusetts, USA). All other chemicals were of analytical grade and purchased from Honeywell Fluka (Loughborough, UK), Mallinckrodt Baker (Phillipsburg, NJ, USA), Merck (Darmstadt, Germany), and Sigma-Aldrich (St. Louis, MO, United States).

### Aerobic incubation of strain B50 with sex steroids

The wild-type and the gene-disrupted mutants of strain B50 were used in resting cell biotransformation assays. Bacteria were first aerobically grown in LB broth (1 L in a 2*-*L Erlenmeyer flask) containing E1 (50 μM) as an inducer at 30°C with continuous shaking (150 rpm). Cells were collected through centrifugation (8,000 × *g*, 20 min, 15°C). The cell pellet was resuspended in a chemically defined mineral medium contained NH_4_Cl (2.0 g/L), KH_2_PO_4_ (0.67 g/L), K_2_HPO_4_ (3.95 g/L), MgSO_4_ (2 mM), CaCl_2_ (0.7 mM), filtered vitamin mixture (as described in DSMZ 1116 medium), ethylenediaminetetraacetic acid (EDTA)-chelated mixture of trace elements (Rabus and Widdel, 1995), and sodium selenite (4 μg/L). The cell suspension (OD_600_ = 1; 1 L) was incubated with E1 or other sex steroids (1 mM) and was aerobically incubated at 30°C with continuous shaking (150 rpm) for 24 h. The resulting samples were acidified using 6N HCl and extracted twice by using equal volumes of ethyl acetate. Ethyl acetate fractions were evaporated, and pellets containing estrogen metabolites were stored at - 20°C until further analysis.

### Thin-layer chromatography

Estrogen metabolites were separated on silica gel aluminum thin-layer chromatography (TLC) plates (Silica gel 60 F_254_: thickness, 0.2 mm; 20 × 20 cm; Merck, Darmstadt, Hesse, Germany). Dichloromethane:ethyl acetate:ethanol (7:2:0.025, vol/vol/vol) was used as the developing solvent system for the separation of estrogen metabolites. Compounds were visualized under ultraviolet light at 254 nm and 302 nm or by spraying TLC plates with 30% (vol/vol) H_2_SO_4_.

### HPLC

A reversed-phase HPLC system (Hitachi, Tokyo, Japan) was used for the separation of estrogen metabolites. Separation was achieved on an analytical RP-C_18_ column [Luna C_18_(2), 5 μm, 150 x 4.6 mm; Phenomenex, Torrance, CA, United States] with a flow rate of 0.8 mL/min. Separation was isocratically performed with 30% (vol/vol) acetonitrile containing formic acid (0.1%; vol/vol) serving as the eluent. Steroid products were detected in the range of 200–450 nm by using a photodiode array detector.

### Solid phase extraction

HPLC-purified samples were further concentrated through solid phase extraction (SPE). The 1-mL BAKERBOND™ SPE Octadecyl (C_18_) Disposable Extraction Columns (J.T. Baker, Avantor, Radnor, PA, United States) was preconditioned with 1 mL of methanol and 1 mL of ddH_2_O (pH 2). Samples were evaporated for 30 min to eliminate acetonitrile before loading onto SPE cartridges. Samples were eluted with 1.5 mL of methanol. The eluate was evaporated for further UPLC–HRMS and NMR analysis.

### UPLC–APCI–HRMS

Ethyl acetate extractable samples or HPLC-purified estrogenic metabolites were analyzed on an UPLC system coupled to an APCI–mass spectrometer. Separation was achieved on a reversed-phase C_18_ column (Acquity UPLC® BEH C_18_; 1.7 μm; 100 × 2.1 mm; Waters, Milford, Massachusetts, United States) with a flow rate of 0.4 mL/min at 50°C (column oven temperature). The mobile phase comprised a mixture of two solvents: Solvent A [2% (vol/vol) acetonitrile containing 0.1% (vol/vol) formic acid] and Solvent B [methanol containing 0.1% (vol/vol) formic acid]. Separation was achieved using a linear gradient of Solvent B from 5% to 99% across 12 min. Mass spectrum data were obtained using a Thermo Fisher Scientific Orbitrap Elite Hybrid Ion Trap-Orbitrap Mass Spectrometer (Waltham, MA, United States) equipped with a standard APCI source operating in the positive ion mode. In APCI–HRMS analysis, the capillary and APCI vaporizer temperatures were 120°C and 400°C, respectively; the sheath, auxiliary, and sweep gas flow rates were 40, 5, and 2 arbitrary units, respectively. The source voltage was 6 kV and the current was 15 μA. The parent scan was in the range of *m/z* 50–600. The predicted elemental composition of individual intermediates was calculated using Xcalibur™ Software Mass Spectrometry Software (Thermo Fisher Scientific; Waltham, MA, United States).

### UPLC–ESI–HRMS

HPLC-purified estrogenic metabolites were also analyzed through UPLC–ESI–HRMS on a UPLC system coupled to an ESI–mass spectrometer. UPLC separation was achieved as described above. Mass spectral data were collected in a +ESI mode in separate runs on a Thermo Fisher Scientific Orbitrap Elite Hybrid Ion Trap-Orbitrap Mass Spectrometer (Waltham, MA, United States) operated in a scan mode of 50–500 *m*/*z*. The source voltage was set at 3.2 kV; the capillary and source heater temperatures were 360 °C and 350 °C, respectively; the sheath, auxiliary, and sweep gas flow rates were 30, 15 and 2 arbitary units, respectively. The predicted elemental composition of individual intermediates was calculated using Xcalibur™ Software Mass Spectrometry Software (Thermo Fisher Scientific).

### NMR spectroscopy

^1^H-, ^13^C- and two-dimensional NMR spectra including COSY, HSQC, and HMBC were recorded at 298°K by using an Agilent 600 MHz DD2 spectrometer (Santa Clara, United States). Chemical shifts (δ) were recorded and presented in parts per million with deuterated methanol (99.8%) as the solvent and internal reference.

### PCR

Bacterial genomic DNA (gDNA) was extracted using the Presto Mini gDNA Bacteria Kit (Geneaid, New Taipei City, Taiwan). DNA fragments used for plasmid assembly were amplified with Platinum polymerase (Thermo Fisher Scientific, Waltham, MA, United States), and PCR products were purified using the GenepHlow Gel/PCR Kit (Geneaid, New Taipei City, Taiwan). For colony PCR reactions, a single colony of strain B50 was dissolved in 50 μL of InstaGene™ Matrix (Bio-Rad, Hercules, CA, United States). Strain B50 cell lysate (5 μL) was added into the PCR mixture (25 μL) containing nuclease-free H_2_O, 2× PCR buffer (12.5 μL), dNTPs (0.4 mM), *Taq* polymerase (2.5 U), Red dye and PCR stabilizer (TOOLS, Taipei, Taiwan), and forward and reverse primers (each 400 nM). PCR products were visualized using standard TAE-agarose gel (1%) electrophoresis with SYBR Green I nucleic acid gel stain (Thermo Fisher Scientific, Waltham, MA, USA).

### The fadD3, aedF, and aedK disruption in strain B50

The disruption of individual thiolase genes (*aedF* or *aedF*) or *fadD3* in strain B50 was performed using homologous recombination by a pK18-Cm^R^-pheS** plasmid, as described previously (Hsiao et al. 2021). Briefly, recombinant sites including the 900 base pair upstream/downstream flanking region and the 100 base pair coding sequence of the target genes were cloned into pK18-Cm^R^-pheS** plasmid backbone by using an In-Fusion® HD Cloning Kit (TAKARA Bio; Kusatsu, Shiga, Japan) to generate the plasmid (*aedF*-, *aedK*- or *fadD3*-pK18-Cm^R^-pheS**). This plasmid was electroporated into *E. coli* strain S17-1 using a Gene Pulser Xcell^™^ (Bio-Rad, Hercules, CA, United States) with the conditions of 2.5kV, 25μF, and 200Ω. The transformed *E. coli* strain S17-1 was co-incubated with wild-type *Rhodococcus* sp. strain B50 at 30°C overnight for horizontal gene transfer through conjugation. Successfully transformed colonies of *Rhodococcus* sp. strain B50 were selected with nalidixic acid and chloramphenicol. The insertion of the chloramphenicol-resistant gene into the strain B50 genome was confirmed using the plasmid-specific primer pairs 5′-TTCATCATGCCGTTTGTGAT-3′ (Pk18-cmpheS-F) and 5′-ATCGTCAGACCCTTGTCCAC-3′ (Pk18-cmpheS-R). The genotypes of the *aedF-*, *aedK*-, and *fadD3*-disrupted mutants were examined using the gene-specific primer pairs: 5′-GCGTCACCCGGATCTGAAGA-3′ (*aedF*-3k-F) and 5′-GTCGGTTGAATTGGACGAGTGTG-3′ (*aedF*-3k-R), 5′-CCGCGAAACATCTTCCTC-3′ (*aedK*-3k-F) and 5′-CCGCCGCCATCCCGTAGG-3′ (*aedK*-3k-R), and 5′-GATTCTCTTCGAGCCACTGC-3′ (*fadD3*-3k-F) and 5′-GCAGATCCACTACTTCGCTC-3′ (*fadD3*-3k-R), respectively (**Table S1**).

### RNA extraction of strain B50

Strain B50 was incubated in the chemically defined mineral medium supplemented with 1 mM E1, testosterone, or cholesterol as the sole carbon source. Bacterial cells were harvested when the substrate was consumed. Cell pellets were resuspended in 200 μL of lysozyme buffer containing 20 mM Tris-HCl, 2 mM EDTA, 1% Triton X-100, and 0.8 mg of lysozyme. After incubation at 37°C for 1 h, 600 μL of TRI reagent (Sigma-Aldrich, St. Louis, MO, United States) was added into the sample, which was maintained at −80°C until further RNA extraction. Total RNA was extracted using the Direct-zol RNA MiniPrep kit (Zymo Research, Irvine, CA, United States), and residual DNA was removed using the TURBO DNA-free kit (Thermo Fisher Scientific, Waltham, MA, United States). The quality and quantity of resulting RNA samples were determined using BioAnalyzer 2100 (Agilent, Santa Clara, CA, United States) and Qubit RNA HS assay kit (Thermo Fisher Scientific, Waltham, MA, United States), respectively, whereas the DNA-removing efficacy was evaluated using the Qubit 1× dsDNA HS assay kit (Thermo Fisher Scientific, Waltham, MA, United States).

### RNA-Seq sequencing of strain B50

Ribosomal RNA was removed from strain B50 transcriptomes by using the Ribo-Zero Magnetic Kit (EPICENTRE Biotechnologies, Madison, WI, United States). Further library construction, including cDNA synthesis, adaptor ligation, and enrichment, was performed according to the instruction of the TruSeq Stranded mRNA Library Prep (Illumina, San Diego, CA, United States). The constructed libraries were sequenced as pair-end reads (with a 301-bp read length) on the Illumina MiSeq system (Illumina, San Diego, CA, United States). The strain B50 genome (accession no.: WPAG00000000.1) and transcriptomes were uploaded onto KBase (Arkin *et al*., 2018). Detailed information and datasets are accessible online (https://kbase.us/n/89341/15/), and the protocol for the transcriptomic analysis is described as follows: **Step 1**, the strain 50 genome was uploaded and used as a reference genome; **step 2**, the three transcriptomes (pair-end reads in the FASTQ format) were uploaded using the “Import FASTQ/SRA File as Reads from Staging Area” application; **step 3**, the uploaded transcriptomic data were grouped into a RNA-seq dataset by using the “Create RNA-seq Sample Set” application; **step 4**, forward and reverse reads were merged and trimmed using the “Trim Reads with Trimmomatic-v0.36” application under the default setting; **step 5**, this trimmed set was aligned to reference genomes by using HISAT2 and the corresponding application was “Align reads using HISAT2-V2.1.0”; and **step 6**, StringTie was used to assemble RNA-seq reads by using the “Assemble Transcripts using StingTie-v1.3.3b” application. Finally, gene expression under different growth conditions (growth with E1, testosterone, or cholesterol) is shown as fragments per kilobase of transcript per million values in **Dataset S1**.

## Acknowledgments

This study was supported by the Ministry of Science and Technology of Taiwan (109-2221-E-001-002, 109-2811-B-001-513, and 110-2311-B-031-001). Po-Hsiang Wang was supported by the Research and Development Office as well as Research Center for Sustainable Environmental Technology, National Central University, Taiwan. We thank the Institute of Plant and Microbial Biology, Academia Sinica, for providing access to the Small Molecule Metabolomics Core Facility (for UPLC–HRMS analyses). We also thank the NGS High Throughput Genomics Core at BRCAS, Academia Sinica for MiSeq sequencing.

## Author Contributions

Y.-R.C. and Y.-L.C. designed the study. T.-H.H., T.-H.L., and M.-R.C. performed the experiments. M.M., M.H., and T.H. contributed new reagents and analytic tools. Y.-L.C., T.-H.H., and Y.-R.C analyzed data. Y.-R.C and P.-H.W. drafted the manuscript. All authors reviewed the manuscript.

## References

Arkin, A.P., Cottingham, R.W., Henry, C.S., Harris, N.L., Stevens, R.L., Maslov, S., et al. (2018) KBase: The United States Department of Energy Systems Biology Knowledgebase. Nat Biotechnol 36: 566–569

Balbi, T., Ciacci, C., and Canesi, L. (2019) Estrogenic compounds as exogenous modulators of physiological functions in molluscs: Signaling pathways and biological responses. Comp Biochem Physiol C Toxicol Pharmacol 222: 135–144.

Baronti, C., Curini, R., D’Ascenzo, G., Di Corcia, A., Gentili, A., and Samperi, R. (2000) Monitoring Natural and Synthetic Estrogens at Activated Sludge Sewage Treatment Plants and in a Receiving River Water. Environ Sci Technol 34: 5059–5066.

Belfroid, A.C., Van der Horst, A., Vethaak, A.D., Schäfer, A.J., Rijs, G.B., Wegener, J., and Cofino, W.P. (1999) Analysis and occurrence of estrogenic hormones and their glucuronides in surface water and waste water in The Netherlands. Sci Total Environ 225: 101–108.

Bergstrand, L.H., Cardenas, E., Holert, J., Van Hamme, J.D., and Mohn, W.W. (2016) Delineation of steroid-degrading microorganisms through comparative genomic analysis. mBio 7: e00166.

Chen, Y.-L., Fu, H.-Y., Lee, T.-H., Shih, C.-J., Huang, L., Wang, Y.-S., et al. (2018) Estrogen Degraders and Estrogen Degradation Pathway Identified in an Activated Sludge. Appl Environ Microbiol 84: e00001–18.

Chen, Y.L., Yu, C.P., Lee, T.H., Goh, K.S., Chu, K.H., Wang, P.H., et al. (2017) Biochemical Mechanisms and Catabolic Enzymes Involved in Bacterial Estrogen Degradation Pathways. Cell Chem Biol 24: 712–724.

Chiang, Y.R., Wei, S.T.S., Wang, P.H., Wu, P.H., and Yu, C.P. (2020) Microbial degradation of steroid sex hormones: implications for environmental and ecological studies. Microb Biotechnol 13: 926–949.

Coombre, R.G., Tsong, Y.Y., Hamilton, P.B., and Sih, C.J. (1966) Mechanisms of steroid oxidation by microorganisms. X. Oxidative cleavage of estrone. J Biol Chem 241: 1587–1595.

Crowe, A.M., Workman, S.D., Watanabe, N., Worrall, L.J., Strynadka, N.C.J., and Eltis, L.D. (2018) IpdAB, a virulence factor in *Mycobacterium tuberculosis*, is a cholesterol ring-cleaving hydrolase. Proc Natl Acad Sci USA 115: E3378–E3387.

Fujii, K., Kikuchi, S., Satomi, M., Ushio-Sata, N., and Morita, N. (2002) Degradation of 17β-Estradiol by a Gram-Negative Bacterium Isolated from Activated Sludge in a Sewage Treatment Plant in Tokyo, Japan. Appl Environ Microbiol 68: 2057–2060.

Hamid, H., and Eskicioglu, C. (2012) Fate of estrogenic hormones in wastewater and sludge treatment: A review of properties and analytical detection techniques in sludge matrix. Water Res 46: 5813–5833.

Hanselman, T.A., Graetz, D.A., and Wilkie, A.C. (2003) Manure-borne estrogens as potential environmental contaminants: a review. Environ Sci Technol 37: 5471–5478.

Harvey, R.A. and Ferrier, D.R. (2011) Biochemistry. Lippincott Williams & Wilkins. Baltimore, MD.

Holert, J., Yücel, O., Jagmann, N., Prestel, A., Möller, H.M., and Philipp, B. (2016) Identification of bypass reactions leading to the formation of one central steroid degradation intermediate in metabolism of different bile salts in *Pseudomonas* sp. strain Chol1. Environ Microbiol 18: 3373–3389.

Huang, C.-H., and Sedlak, D.L. (2001) Analysis of estrogenic hormones in municipal wastewater effluent and surface water using enzyme-linked immunosorbent assay and gas chromatography/tandem mass spectrometry. Environ Toxicol Chem 20: 133–139.

Hsiao, T.H., Chen, Y.L., Meng, M., Chuang, M.R., Horinouchi, M., Hayashi, T., Wang, P.H., and Chiang, Y.R. (2021) Mechanistic and phylogenetic insights into actinobacteria-mediated estrogen biodegradation in urban estuarine sediments. Microb Biotechnol doi: 10.1111/1751-7915.13798.

Ibero, J., Galán, B., Díaz, E., and García, J.L. (2019a) Testosterone degradative pathway of *Novosphingobium tardaugens*. Genes 10: 871.

Ibero, J., Sanz, D., Galán, B., Díaz, E., and García, J.L. (2019b) High-quality whole-genome sequence of an estradiol-degrading strain, *Novosphingobium tardaugens* NBRC 16725. Microb Resour Annouc 8: e01715–e01718.

Ibero, J., Galán, B., Rivero-Buceta, V., García, J.L. (2020) Unraveling the 17β-estradiol degradation pathway in *Novosphingobium tardaugens* NBRC 16725. Front Microbiol 11: 588300.

Jurgens, M.D., Holthaus, K.I.E., Johnson, A.C., Smith, J.J.L., Hetheridge, M., and Williams, R.J. (2002) The potential for estradiol and ethinylestradiol degradation in English Rivers. Environ Toxicol Chem 21: 480–488.

Ke, J., Zhuang, W., Gin, K.Y.H., Reinhard, M., Hoon, L.T., and Tay, J.H. (2007) Characterization of estrogen-degrading bacteria isolated from an artificial sandy aquifer with ultrafiltered secondary effluent as the medium. Appl Microbiol Biotechnol 75: 1163–1171.

Koh, Y.K.K., Chiu, T.Y., Boobis, A., Cartmell, E., Scrimshawm, M.D., and Lester, J.N. (2008) Treatment and removal strategies for estrogens from wastewater. Environ Technol 29: 245–267

Kolodziej, E.P., Gray, J.L., and Sedlak, D.L. (2003) Quantification of steroid hormones with pheromonal properties in municipal wastewater effluent. Environ Toxicol Chem 22: 2622–2629.

Kolodziej, E.P., Harter, T., and Sedlak, D.L. (2004) Dairy wastewater, aquaculture, and spawning fish as sources of steroid hormones in the aquatic environment. Environ Sci Technol 38: 6377–6384.

Kramer, V.J., Miles-Richardson S., Pierens S.L., and Giesy J.P. (1998) Reproductive impairment and induction of alkaline-labile phosphate, a biomarker of estrogen exposure, in fathead minnows (*Pimephales promelas*) exposed to waterborne 17β-estradiol. Aquat Toxicol 40: 335–360.

Kurisu, F., Ogura, M., Saitoh, S., Yamazoe, A., and Yagi, O. (2010) Degradation of natural estrogen and identification of the metabolites produced by soil isolates of *Rhodococcus* sp. and *Sphingomonas* sp. J Biosci Bioeng 109: 576–582.

Lee, Y.C., Wang, L.M., Xue, Y.H., Ge, N.C., Yang, X.M., and Chen, G.H. (2006) Natural Estrogens in the Surface Water of Shenzhen and the Sewage Discharge of Hong Kong. Hum Ecol Risk Assess Int J 12: 301–312.

Li, S., Sun, K., Yan, X., Lu, C., Waigi, M.G., Liu, J., and Ling, W. (2021) Identification of novel catabolic genes involved in 17β-estradiol degradation by *Novosphingobium* sp. ES2-1. Environ Microbiol doi: 10.1111/1462-2920.15475.

Lin, A.Y., and Reinhard, M. (2005) Photodegradation of common environmental pharmaceuticals and estrogens in river water. Environ Toxicol Chem 24: 1303–1309.

Lin, C.W., Wang, P.H., Ismail, W., Tsai, Y.W., El Nayal, A., Yang, C.Y., et al. (2015) Substrate uptake and subcellular compartmentation of anoxic cholesterol catabolism in *Sterolibacterium denitrificans*. J Biol Chem 290: 1155–1169.

Lorenzen, A., Hendel, J.G., Conn, K.L., Bittman, S., Kwabiah, A.B., Lazarovitz, G., et al. (2004) Survey of hormone activities in municipal biosolids and animal manures. Environ Toxicol 19: 216–225.

Matsumoto, T., Osada, M., Osawa, Y., and Mori, K. (1997) Gonadal Estrogen Profile and Immunohistochemical Localization of Steroidogenic Enzymes in the Oyster and Scallop during Sexual Maturation. Comp Biochem Physiol B: Biochem Mol Biol 118: 811–817.

Noguera-Oviedo, K., and Aga, D.S. (2016) Chemical and biological assessment of endocrine disrupting chemicals in a full scale dairy manure anaerobic digester with thermal pretreatment. Sci Total Environ 550: 827–834.

Pelletier, D.A., and Harwood, C.S. (1998) 2-Ketocyclohexanecarboxyl coenzyme A hydrolase, the ring cleavage enzyme required for anaerobic benzoate degradation by *Rhodopseudomonas palustris*. J Bacteriol 180: 2330–2336.

Pelletier, D.A., Harwood, C.S. (2000) 2-Hydroxycyclohexanecarboxyl coenzyme A dehydrogenase, an enzyme characteristic of the anaerobic benzoate degradation pathway Used by *Rhodopseudomonas palustris*. J Bacteriol 182: 2753–2760.

Qin, D., Ma, C., Lv, M., and Yu, C.P. (2020) *Sphingobium estronivorans* sp. nov. and *Sphingobium bisphenolivorans* sp. nov., isolated from a wastewater treatment plant. Int J Syst Evol Microbiol 70: 1822–1829.

Rabus, R., and Widdel, F. (1995) Anaerobic degradation of ethylbenzene and other aromatic hydrocarbons by new denitrifying bacteria. Arch Microbiol 163: 96–103.

Takamura, Y., and Nomura, G. (1988) Changes in the intracellular concentration of acetyl-CoA and malonyl-CoA in relation to the carbon and energy metabolism of *Escherichia coli* K12. J Gen Microbiol 134: 2249–2253.

Tarrant, A.M., Blomquist, C.H., Lima, P.H., Atkinson, M.J., and Atkinson, S. (2003) Metabolism of estrogens and androgens by scleractinian corals. Comp Biochem Physiol B Biochem Mol Biol 136: 473–485.

Thayanukul, P., Zang, K., Janhom, T., Kurisu, F., Kasuga, I., and Furumai, H. (2010) Concentration-dependent response of estrone-degrading bacterial community in activated sludge analyzed by microautoradiography-fluorescence in situ hybridization. Water Res 44: 4878–4887.

Wang, P.H., Chen, Y.L., Wei, S.T.S., Wu, K., Lee, T.H., Wu, T.Y., and Chiang, Y.R. (2020) Retroconversion of estrogens into androgens by bacteria via a cobalamin-mediated methylation. Proc Natl Acad Sci USA 117: 1395–1403.

Wu, K., Lee, T.H., Chen, Y.L., Wang, Y.S., Wang, P.H., Yu, C.P., et al. (2019) Metabolites Involved in Aerobic Degradation of the A and B Rings of Estrogen. Appl Environ Microbiol 85: e02223–18.

Yoshimoto, T., Nagai, F., Fujimoto, J., Watanabe, K., Mizukoshi, H., Makino, T., et al. (2004) Degradation of estrogens by Rhodococcus zopfii and Rhodococcus equi isolates from activated sludge in wastewater treatment plants. Appl Environ Microbiol 70: 5283–5289.

Yu, C.P., Deeb, R.A., and Chu, K.H. (2013) Microbial degradation of steroidal estrogens. Chemosphere 91: 1225–1235.

Yu, C.P., Roh, H., and Chu, K.H. (2007) 17β-Estradiol-Degrading Bacteria Isolated from Activated Sludge. Environ Sci Technol 41: 486–492.

Zhao, H., Tian, K., Qiu, Q., Wang, Y., Zhang, H., Ma, S., Jin, S., and Huo, H. (2018) Genome analysis of *Rhodococcus* Sp. DSSKP-R-001: A highly effective β-estradiol-degrading bacterium. Int J Genomics 2018: 3505428.

