## Supplemental Information for "Identification of essential β-oxidation genes and corresponding metabolites for estrogen degradation by actinobacteria"

Running title: Actinobacterial estrogen degradation pathway

**Tsun-Hsien Hsiao[1], Tzong-Huei Lee [2], Meng-Rong Chuang[1], Po-Hsiang Wang [3,4], Menghsiao Meng[5], Masae Horinouchi[6], Toshiaki Hayashi[7], Yi-Lung Chen[8*], and Yin-Ru Chiang[1*]**

[1]Biodiversity Research Center, Academia Sinica, Taipei 115, Taiwan

[2]Institute of Fisheries Science, National Taiwan University, 106 Taipei, Taiwan

[3]Gradaute Institute of Environmental Engineering, National Central University, Taoyuan 320, Taiwan

[4]Earth-Life Science Institute (ELSI), Tokyo Institute of Technology, Tokyo, Japan

[5]Graduate Institute of Biotechnology, National Chung Hsing University, Taichung 402, Taiwan

[6]Condensed Molecular Materials Laboratory, RIKEN, Saitama, 351-0198 Japan

[7]Environmental Molecular Biology Laboratory, RIKEN, Saitama, 351-0198 Japan

[8]Department of Microbiology, Soochow University, Taipei 111, Taiwan

**Table S1.** Oligonucleotides used in this study.

| Primer | Sequence (5'- 3') | Usage |
|---|---|---|
| *fadD3*-UP-F<br>*fadD3*-UP-R | ACACAGGAAACAGCTATGACACCAAGCAATTGCTAGGTAGGGAG<br>CCGCGACCTCCGGTGAGCACCGCGTCGAAG | For strain B50 *fadD3* up-stream recombinant fragment cloning |
| *fadD3*-DOWN-F<br>*fadD3*-DOWN-R | GTGCTCACCGGAGGTCGCGGTGGTCGGTG<br>GTTGTAAAACGACGGCCAGTGCATGGGCGAAGTGGTGC | For strain B50 *fadD3* down-stream recombinant fragment cloning |
| *aedF*-UP-F<br>*aedF*-UP-R | ACACAGGAAACAGCTATGACGAGGGTGGCGACGGCACC<br>TTCGTGCAGCGTACGTGAGCGCCGAGGTCG | For strain B50 *aedF* up-stream recombinant fragment cloning |
| *aedF*-DOWN-F<br>*aedF*-DOWN-R | GCTCACGTACGCTGCACGAACTCGAACGC<br>GTTGTAAAACGACGGCCAGTATTTCGTCGAGTCGATCGAGC | For strain B50 *aedF* down-stream recombinant fragment cloning |
| *aedK*-UP-F<br>*aedK*-UP-R | ACACAGGAAACAGCTATGACGAACAATCTTGCCGGAGCACTCC<br>CCGCTGCGGTGCCTCGAGTTGCAGCTCG | For strain B50 *aedK* down-stream recombinant fragment cloning |
| *aedK*-DOWN-F<br>*aedK*-DOWN-R | AACTCGAGGCACCGCAGCGGCAGGTCGC<br>GTTGTAAAACGACGGCCAGTGTAGGTGACATGGGGGTCG | For strain B50 *aedK* down-stream recombinant fragment cloning |
| Pk18-M13-F<br>Pk18-M13-R | ACTGGCCGTCGTTTTACAAC<br>GTCATAGCTGTTTCCTGTGTG | For generating the fragment of pK18-Cm$^{R}$-pheS** as the in-fusion backbone |
| *fadD3*-3k-F<br>*fadD3*-3k-R | GATTCTCTTCGAGCCACTGC<br>GCAGATCCACTACTTCGCTC | For checking the disruption of strain B50 *fadD3* |
| *aedF*-3k-F<br>*aedF*-3k-R | GCGTCACCCGGATCTGAAGA<br>GTCGGTTGAATTGGACGAGTGTG | For checking the disruption of strain B50 *aedF* |
| *aedK*-3k-F<br>*aedK*-3k-R | CCGCGAAACATCTTCCTC<br>CCGCCGCCATCCCGTAGG | For checking the disruption of strain B50 *aedK* |
| Pk18-cmpheS-F<br>Pk18-cmpheS-R | TTCATCATGCCGTTTGTGAT<br>ATCGTCAGACCCTTGTCCAC | For checking the insertion of pK18-Cm$^{R}$-pheS** plasmid |

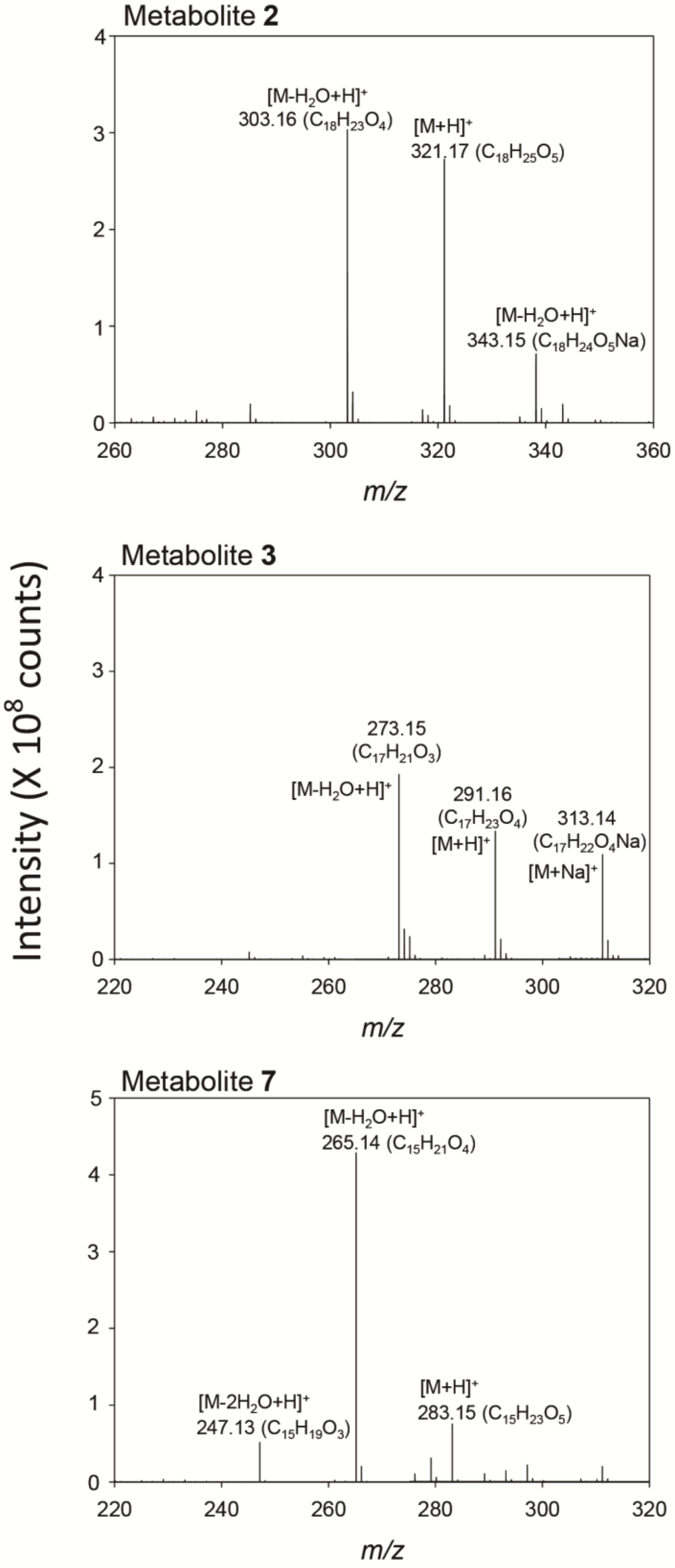


**Figure S1** ESI–HRMS spectra of three novel estrogenic metabolites produced by strain B50.

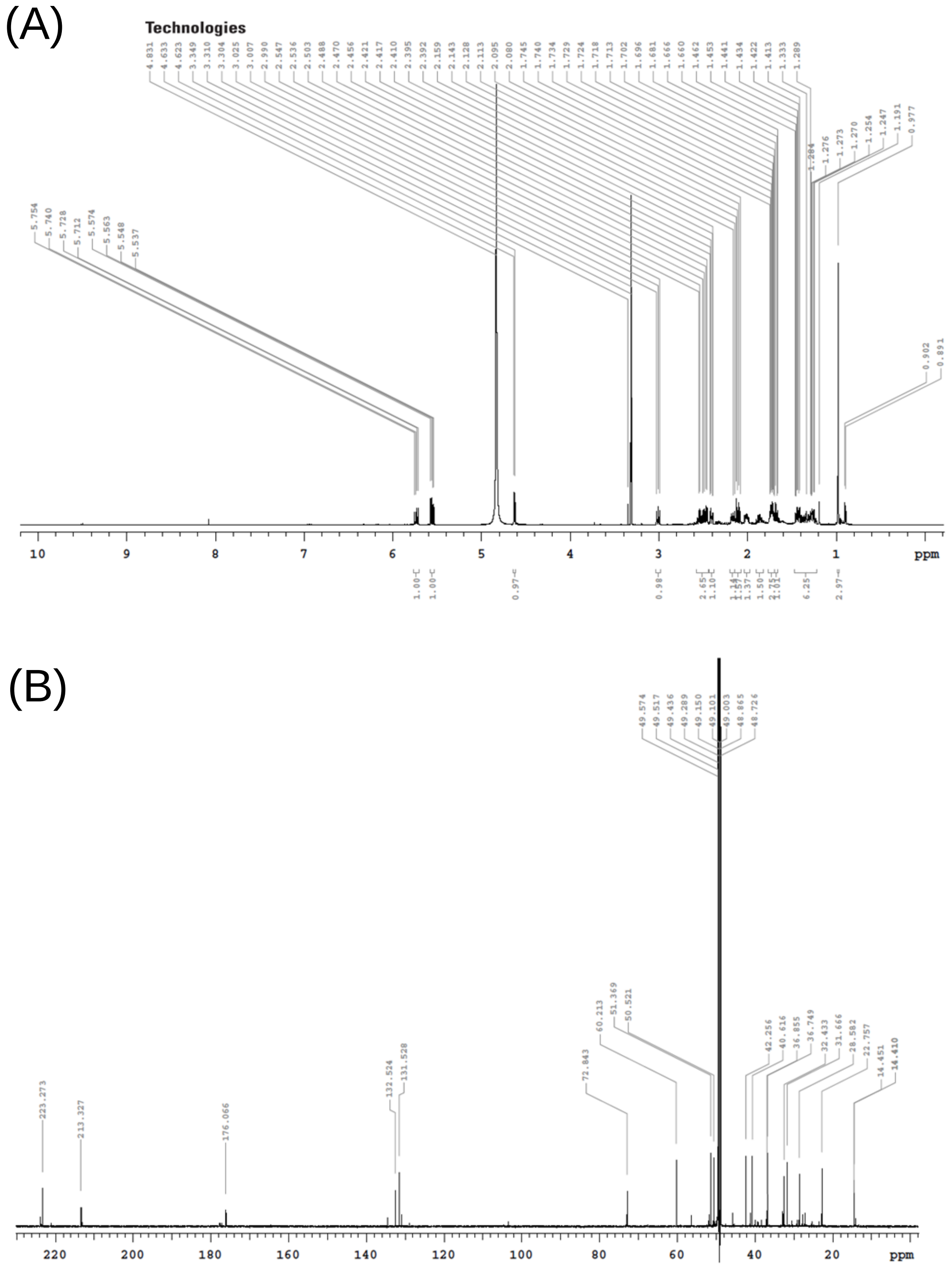


**Figure S2** $^{1}$H- (500 MHz) (A) and $^{13}$C-NMR (125 MHz) (B) spectra of Metabolite **2**.

(A)

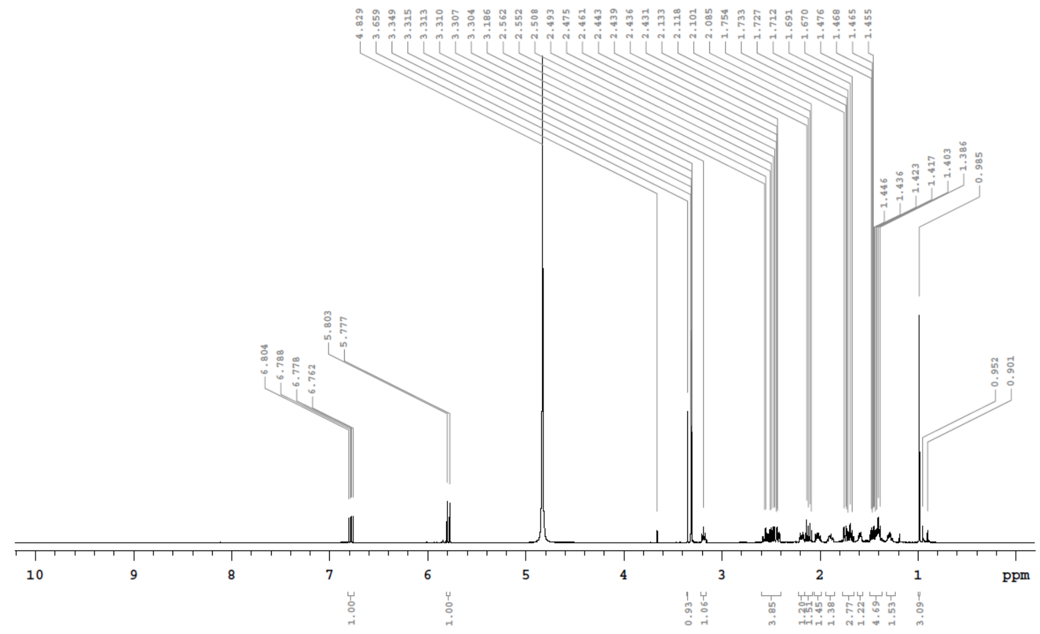


(B)

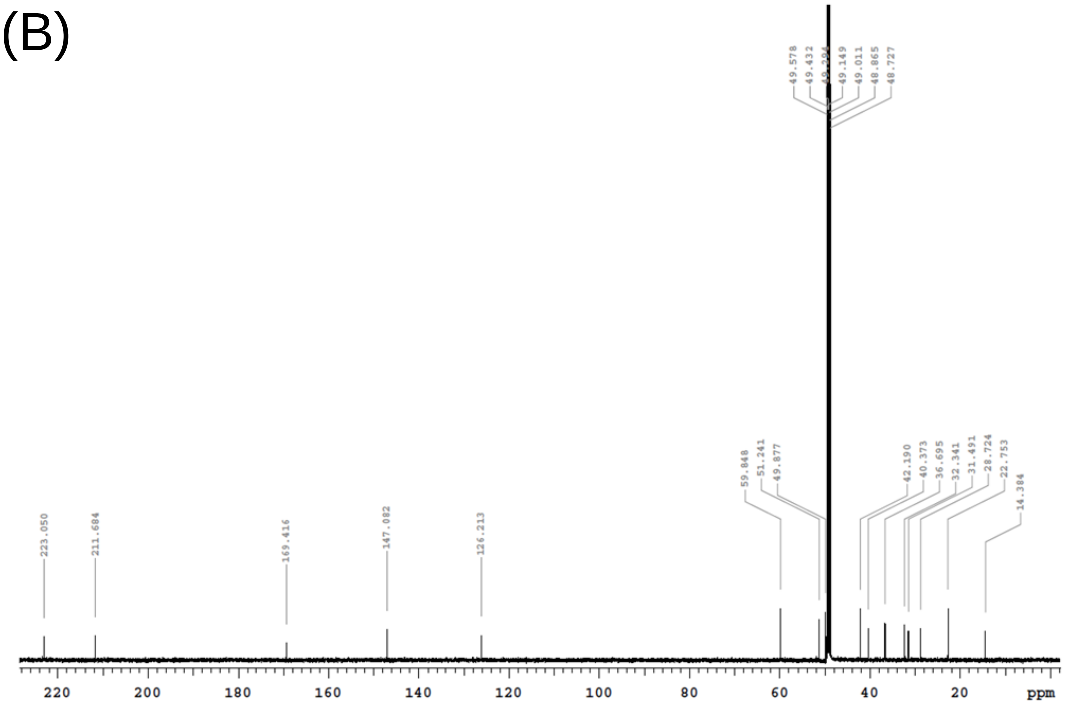


**Figure S3** $^{1}$H- (500 MHz) (A) and $^{13}$C-NMR (125 MHz) (B) spectra of Metabolite **3**.

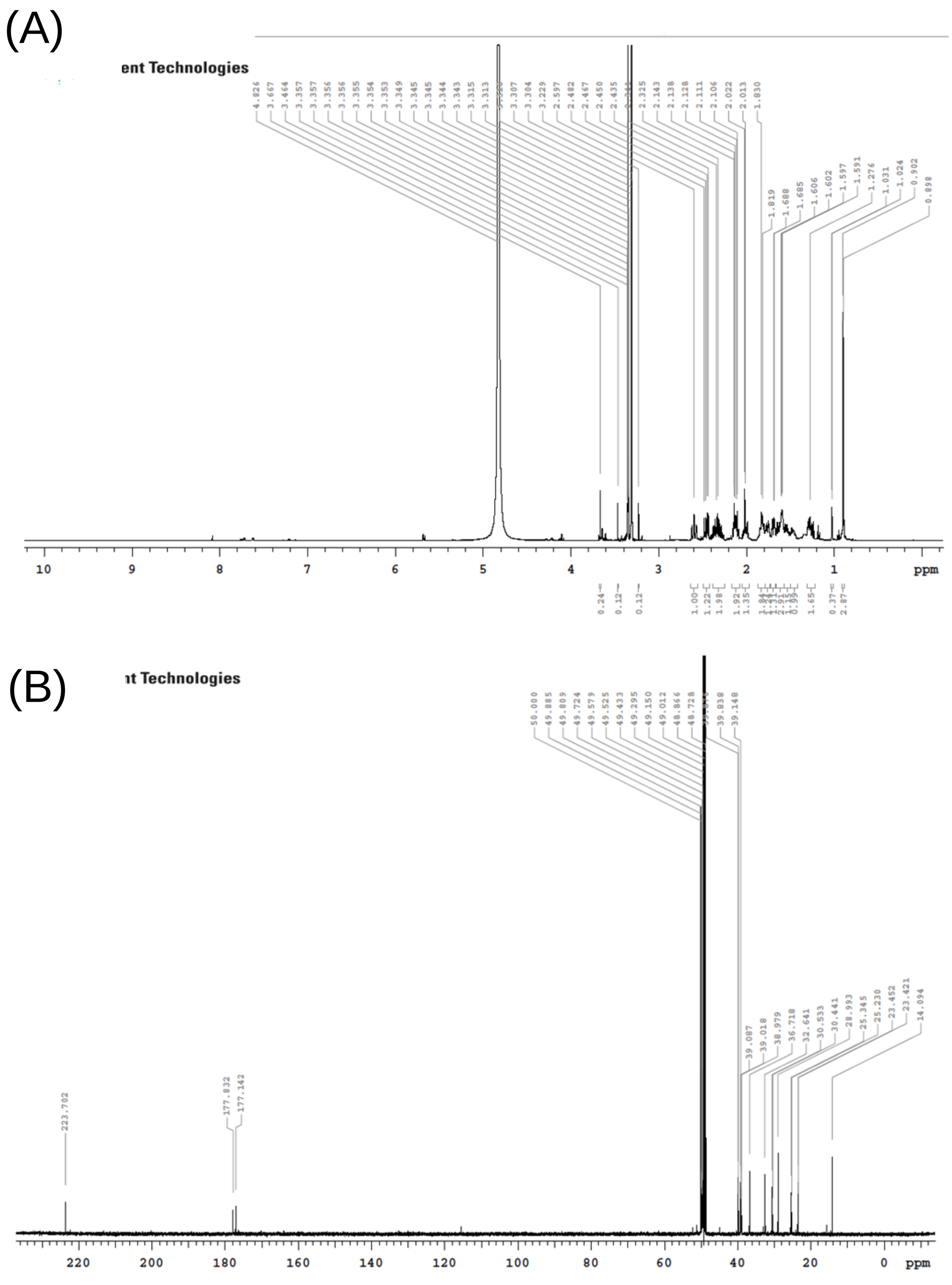


**Figure S4** $^{1}$H- (500 MHz) (A) and $^{13}$C-NMR (125 MHz) (B) spectra of Metabolite **7**.

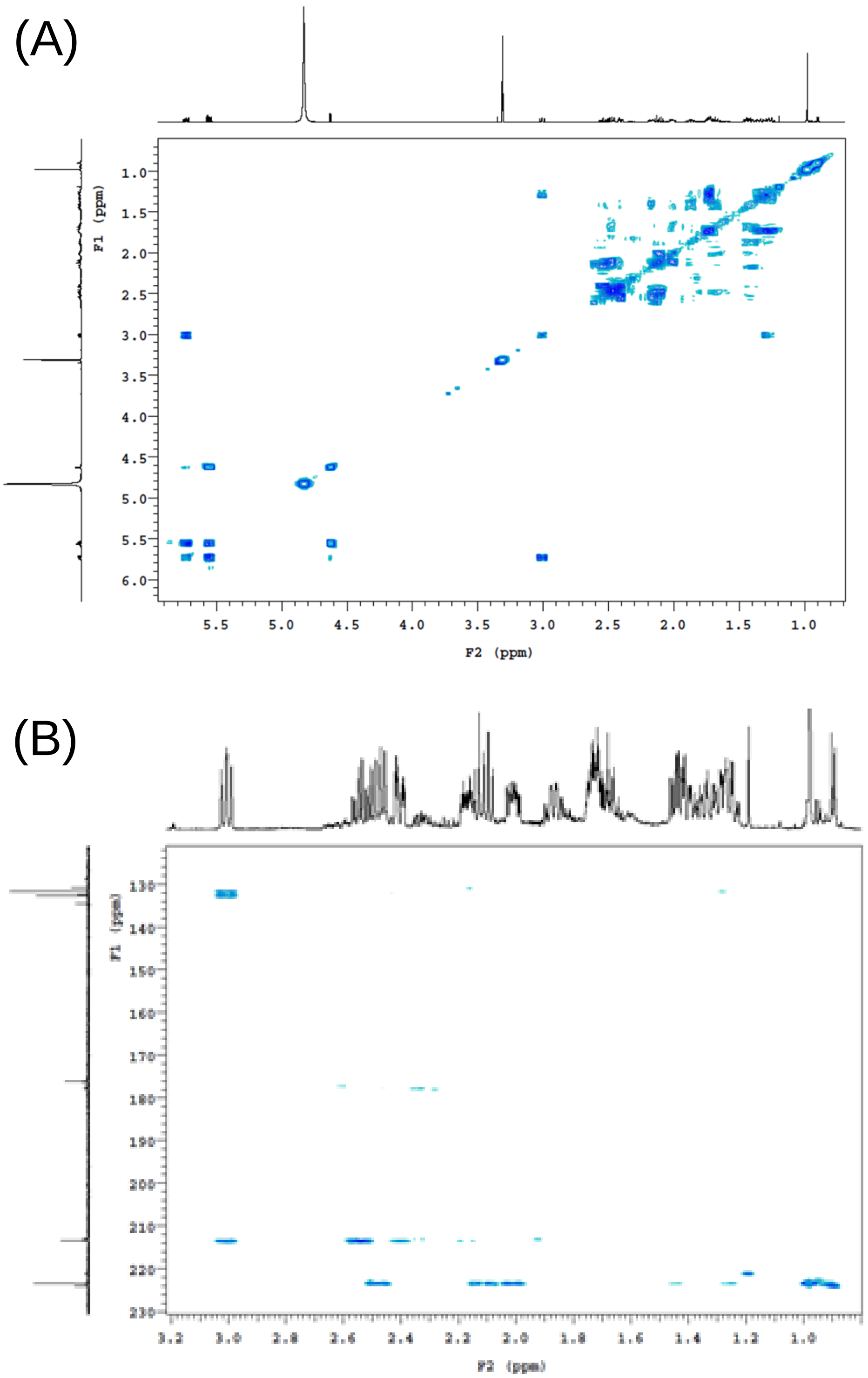


**Figure S5** COSY (A) and HMBC (B) spectra of Metabolite **2**.

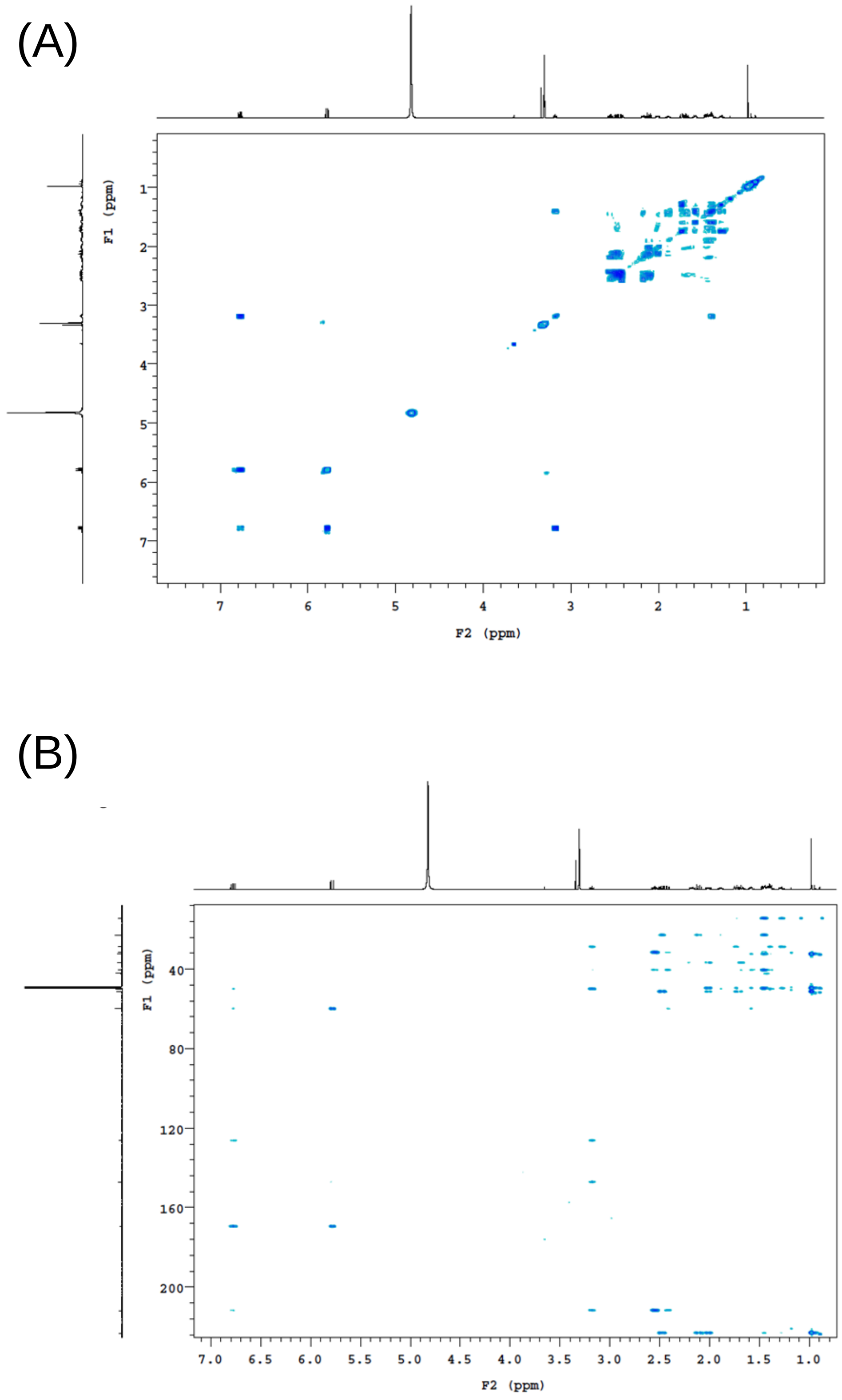


**Figure S6** COSY (A) and HMBC (B) spectra of Metabolite **3**.

(A)

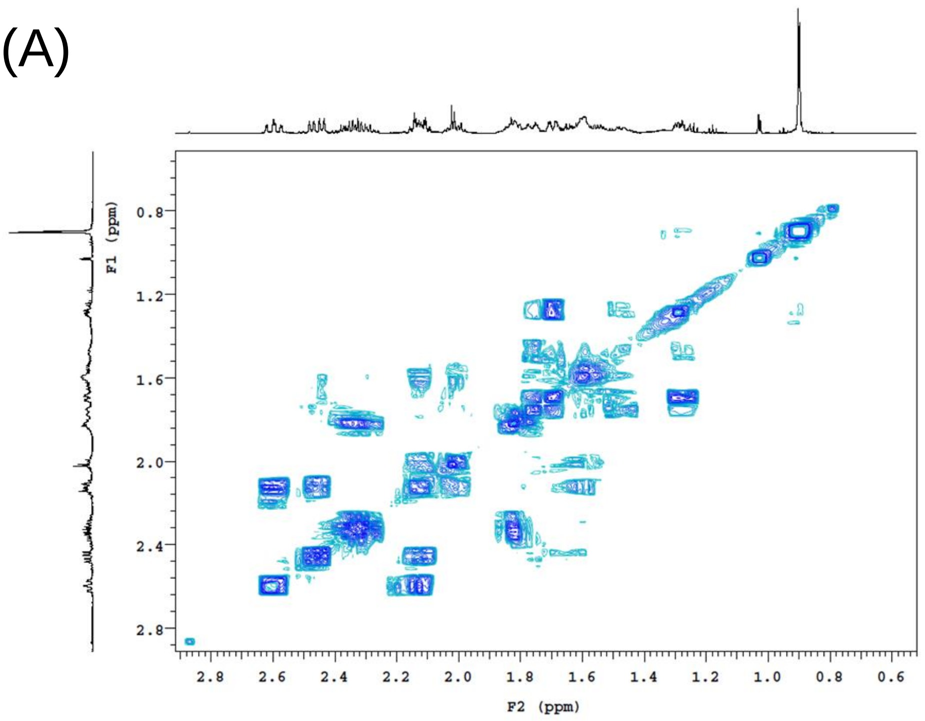


(B)

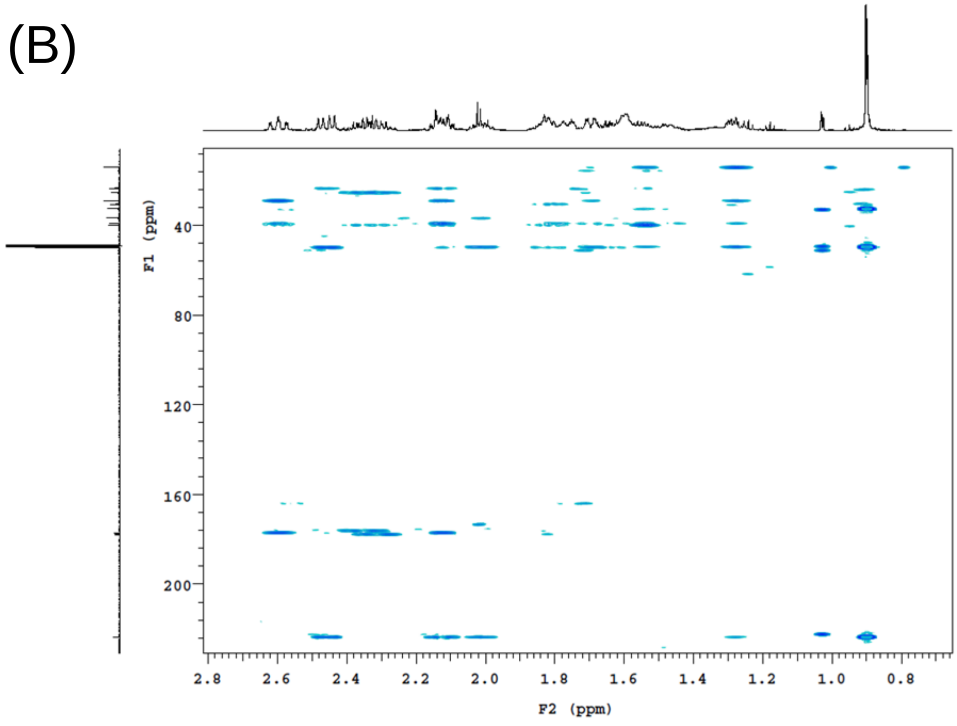


**Figure S7** COSY (A) and HMBC (B) spectra of Metabolite **7**.

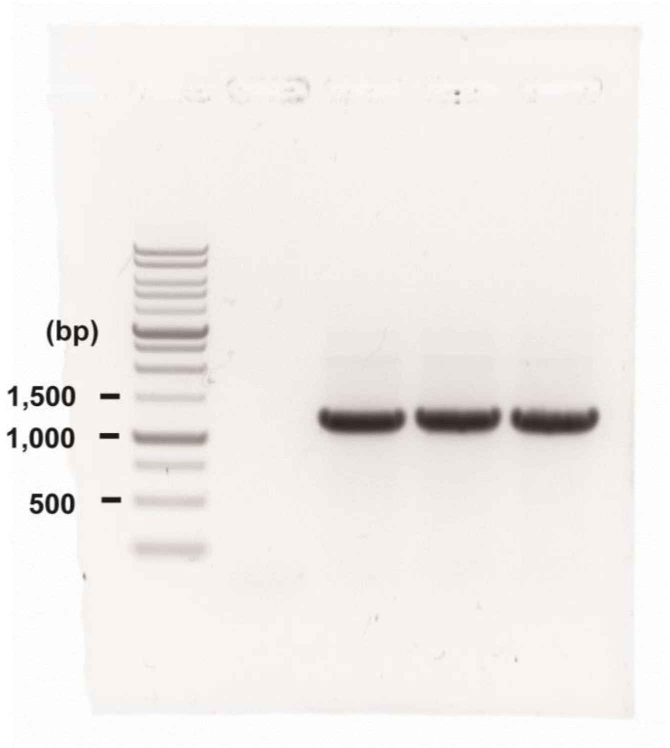


**Figure S8** The uncropped full-range agarose gel (1%) of **Fig. 3Bi.**

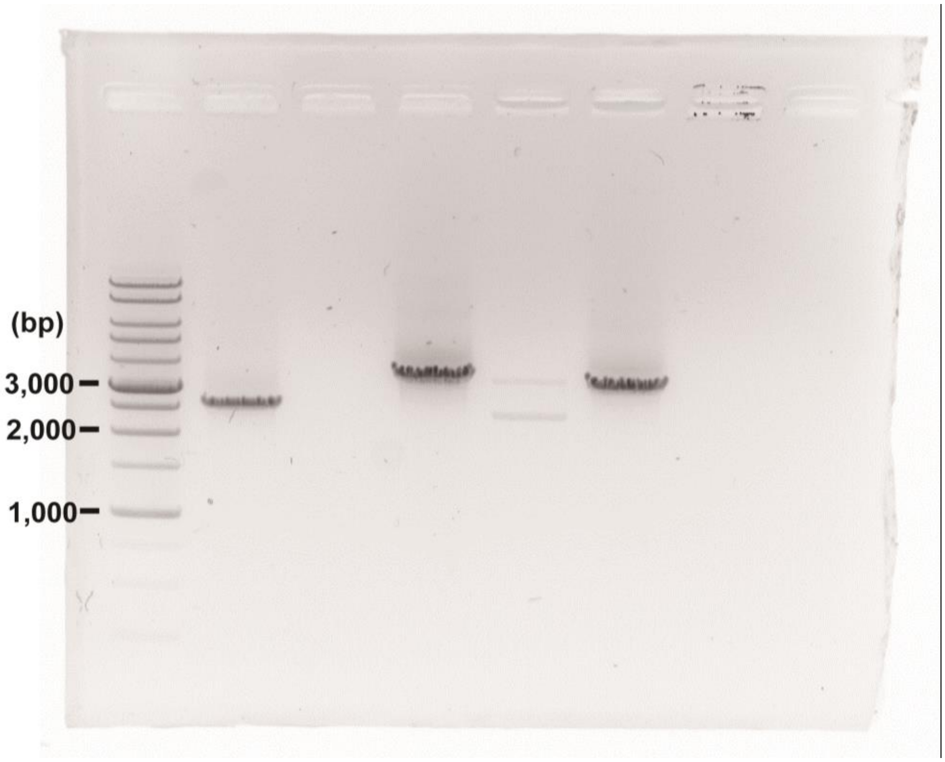


**Figure S9** The uncropped full-range agarose gel (1%) of **Fig. 3Bii**.
